# A Distinct Chromatin State Drives Therapeutic Resistance in Invasive Lobular Breast Cancer

**DOI:** 10.1101/2022.03.29.486217

**Authors:** Agostina Nardone, Xintao Qiu, Ariel Feiglin, Xiaoyong Fu, Sandor Spisak, Avery Feit, Gabriela Cohen Feit, Yingtian Xie, Alba Font-Tello, Cristina Guarducci, Francisco Hermida-Prado, Sudeepa Syamala, Klothilda Lim, Matthew Pun, MacIntosh Cornwell, Weihan Liu, Aysegul Ors, Hisham Mohammed, Jane Brock, Matthew L. Freedman, Rachel Schiff, Eric P. Winer, Henry Long, Otto Metzger Filho, Rinath Jeselsohn

## Abstract

Most invasive lobular breast cancers (ILC) are of the luminal A subtype and strongly hormone receptor positive. Yet, they are relatively resistant to tamoxifen and are associated with inferior long-term outcomes compared to invasive ductal cancers (IDC). In this study, we sought to gain mechanistic insights into these clinical findings that are not explained by the genetic landscape of ILC and to identify strategies to improve patient outcomes. Through a comprehensive analysis of the epigenome of ILC in pre-clinical models and clinical samples we found that compared to IDC, ILC has a distinct chromatin state that is linked to gained recruitment of FOXA1, a lineage-defining pioneer transcription factor. This results in an ILC-unique FOXA1-estrogen receptor (ER) axis that promotes the transcription of genes associated with tumor progression and poor outcomes. The ILC-unique FOXA1-ER axis leads to retained ER chromatin binding after tamoxifen treatment thereby facilitating tamoxifen resistance while remaining strongly dependent on ER signaling. Mechanistically, gained FOXA1 binding was associated with the auto-induction of FOXA1 in ILC through an ILC-unique FOXA1 binding site. Targeted silencing of this regulatory site resulted in the disruption of the feed-forward loop and growth inhibition in ILC.

In summary, we show that ILC is characterized by a unique cell state and FOXA1-ER axis that dictate tumor progression and offer a novel mechanism of tamoxifen resistance. These results underscore the importance of conducting clinical trials dedicated to patients with ILC to optimize endocrine treatments in this breast cancer subtype.

## Introduction

Invasive lobular carcinoma (ILC) is the second most common histological subtype of breast cancer (BC), accounting for 10-15% of all cases. The classical variant of ILC is characterized by relatively uniform and non-cohesive cells that grow as a single file infiltrating the stroma. This growth pattern is attributed to the loss of E-cadherin, the hallmark of ILC (1,2), and can render physical exam and mammographic diagnosis challenging (3,4). Although the majority of ILCs are of low or intermediate grade, express high levels of estrogen receptor (ER), and of luminal A subtype, several studies suggest that ILC long-term outcomes are inferior to stage-matched invasive ductal carcinoma (IDC) (5). Additionally, the retrospective analysis of the BIG-1-98 adjuvant endocrine trial demonstrated that the magnitude of the inferior benefit from tamoxifen compared with an aromatase inhibitor (AI) was greater in ILC compared with IDC (6), indicative of relative resistance to tamoxifen treatment in early-stage treatment naïve ILC.

Several studies have revealed differences in the mutational landscape of ILC compared to IDC. In addition to loss of E-cadherin, ILC is characterized by a higher frequency of FOXA1 mutations (7,8), which are found in 7-9% of primary ILC versus 2% in primary IDC. In primary, treatment naïve, ILC tumors, FOXA1 mutations are associated with increased expression of FOXA1 and unique transcriptional profiles (7). More recently, FOXA1 mutations were shown to be associated with endocrine resistance in metastatic ER-positive (ER+) BC (9). FOXA1 is a pioneer transcription factor (TF) that mediates ER transcriptional activity (10–13). Thus, the increase in FOXA1 mutations in ILC and the disparate patterns of response to the different classes of endocrine treatment suggest that ILC may have a unique ER axis compared to IDC. In support of this notion, previous work has shown that ILC cells are characterized by a unique transcriptional response to estrogen (14). However, the mechanism by which the ER transcriptional axis is altered in ILC models lacking FOXA1 mutations remains elusive. In this study, we performed a comprehensive analysis of the epigenome of ILC in pre-clinical models and clinical samples with the aim to provide mechanistic insights to explain the distinctive responses to estradiol and tamoxifen in ILC versus IDC and to inform us on new potential therapeutic approaches for ILC.

## Materials and Methods

### Cell lines

MCF7, T47D and MDA-MB-134-VI (MDAMB134) cells were purchased from ATCC; SUM44PE (SUM44) cells were a gift by Dr. Stephen Ethier. All the cells were authenticated and regularly tested for mycoplasma contamination. The MCF7 cells were maintained in DMEM and T47D cells in RPMI supplemented with 10% heat-inactivated fetal bovine serum (FBS) and 1% penicillin/streptomycin (P/S). MDA134 cells were maintained in L-15 with 20%FBS and 1% P/S. SUM44 were maintained in Ham’s F-12 supplemented with insulin (5ug/mL), hydrocortisone (1ug/mL), fungizone (2.5ug/mL), transferrin (5ug/mL), T3 (6.6ng/mL), ethanolamin (5mM), NaSe (8.7ng/mL) (all from Sigma-Aldrich, St. Louis, MO), gentamicin (25ug/mL), HEPES (10mM), BSA (0.1%) (all from Thermo Fisher Scientific). For hormone-depleted (HD) conditions, cells were kept in phenol-red free medium supplemented with 10% heat-inactivated charcoal-stripped (CS)-FBS (with the exception of SUM44) (20% for MDA134) and 1% P/S. MCF7, T47D and SUM44 were incubated at 37°C in 5% CO_2_, MDA134 were incubated at 37°C without CO_2_.

### Human tissue studies

The ILC metastatic ascitic fluid was obtained with the patient’s written consent and the approval of the Dana-Farber/Harvard Cancer Center institutional review board (Protocol 93-085). Red blood cells and dead cells were removed by Ficoll (Sigma-Aldrich) followed by selection of cancer epithelial cells using EpCam selective beads (Dynabeads). Cancer epithelial cells were subjected to FOXA1, ER and H3K27ac ChIP-seq.

### Chromatin Immunoprecipitation (ChIP)-Sequencing

ChIP experiments were conducted as described previously (15) and were done in duplicates. MCF7, T47D, MDAMB134 and SUM44 cells were cultured for three days in HD conditions then treated with 10nM estradiol for 45 minutes. For the ChIP experiments with tamoxifen we used 10nM 4-hydroxytamoxifen for 45 minutes. Chromatin from 20 million formaldehyde-fixed cells was sonicated to a size range of 200-300 bp. Solubilized chromatin was immunoprecipitated with a mix of the ER antibodies Ab10 (Thermo Fisher Scientific) and SC-543 (Santa Cruz), a mix of the FOXA1 antibodies ab5089 and ab23738 (Abcam), GATA3 antibody D13C9 (Cell signaling) or H3K27Ac antibody (C15410196, Diagenode). The same antibodies were used in each experiment for all the cell lines. The samples were reversed crosslinked, treated with proteinase K, and DNA was extracted. Libraries were sequenced using 75 bp paired-end reads on the Illumina Nextseq500 at the Dana-Farber Cancer Institute.

### RNA sequencing

Ductal and lobular cell lines were cultured for three days in HD conditions. After washing the cells with PBS, cells were incubated for 12 hours in HD medium with 10nM estradiol, or DMSO treatment. Total RNA was extracted using the QIAGEN RNeasy kit with DNAse digestion. RNA concentrations were measured by nanodrop and quality of RNA was determined by a Bioanalyzer. For all cell line studies, samples were analyzed in at least duplicates. RNA-seq libraries were made using the TruSeq RNA Sample Preparation Kit (Illumina). Samples were sequenced on an Illumina Nextseq500.

### CRISPRi

#### FOXA1 enhancer

Stable dCas9-KRAB expressing MCF7 and SUM44 cell lines were generated by lentiviral transduction using Lenti-dCas9-KRAB-blast (Addgene #89567) transfer vector and pMD2.G (Addgene #12259) and psPAX (Addgene#12259) lentiviral packaging vectors. Transduced cells were selected and maintained in blasticidin (10 μg/mL) selective culture media. Guide RNAs (gRNAs) were designed against peak 1 and non-human targeting control (NTC) gRNA was included and cloned as described previously (detailed protocol available at http://www.broadinstitute.org/rnai/public/resources/protocols). Briefly, for each gRNA complementary single-stranded oligonucleotides were synthetized (Invitrogen), phosphorylated, annealed, and gRNA cassettes were ligated into pXPR_BRD003 gRNA expression vector containing puromycin selection marker. After bacterial transformation (Stbl3 cells, Invitrogen) individual clones were picked and regenerated, and correct gRNA sequence were confirmed by Sanger-sequencing.

Lentiviral particles were generated for each sgRNA experiment by transforming HEK-293T cells with gRNA transfer vectors and lentiviral packaging mix (pMD2.G psPAX). Lentiviral particle containing media were collected and filtered using 0.45 μm pore size syringe filters (Corning) after 48h post transfection and used for treatment of previously plated breast cell lines. Media was changed after 24h and replaced by puromycin (2 μg/mL) containing selective culture medium. Puromycin resistant cells were collected after 7 days for cell of selection for proliferation studies and RNA for cDNA.

The coordinates of the region of interest (P1): chromosome 14 from 38,067,100 to 38,071,000, Human hg19. Working guides were: gP1: caccgAGGAGCTACTAGACCAGTAA and aaacTTACTGGTCTAGTAGCTCCTC NTC: caccGCGACCCAAATGCACCCTTT NTC and aaacAAAGGGTGCATTTGGGTCGC

### CRISPRi control studies

MCF7 cells engineered to express dCAS9-KRAB (details in main methods) and parental MCF7 cells were infected with the pKLV2-U6gRNA5(gGFP)-PGKBFP2AGFP-W plasmid (Addgene plasmid # 67980). For the infection the virus was generated in 293FT cells transfected with PAX2 and pMD2.G plasmids together with the pKLV2-U6gRNA5(gGFP)-PGKBFP2AGFP-W plasmid using the X-tremeGENE reagent (Roche). MCF7 cells (without expression of dCCAS9-KRAB or the pKLV2-U6gRNA5(gGFP)-PGKBFP2AGFP-W plasmid), MCF7 cells expressing the pKLV2-U6gRNA5(gGFP)-PGKBFP2AGFP-W plasmid and cells expressing dCAS9-KRAB and the pKLV2-U6gRNA5(gGFP)-PGKBFP2AGFP-W plasmid were analyzed by flow cytometry for GFP (alexa-Flour 488) and BFP (DAPI).

### Quantification and Statistical analysis

Statistical analyses for cell proliferation studies were performed using two-sided Student’s t-tests, and p-values less than 0.05 were considered statistically significant. Error bars represent the ± SEM. Cell line experiments testing cell proliferation were all performed in triplicates.

Additional methods are in the supplemental file.

### Data availability

The whole-genome sequencing, RNA-seq, ChIP-seq, ATAC-seq, and Hi-ChIP-seq data have been deposited in the Gene Expression Omnibus database under GSE152367.

## Results

### ILC has a unique chromatin state that is tightly linked to FOXA1 recruitment

To study the epigenetic landscape and identify the genome-wide active regulatory elements accessible to TF binding in ILC versus IDC we first conducted transposase-accessible chromatin sequencing (ATAC-seq) in cell line models of ER+ ILC (MDA-MB-134-VI (MDAMB134)) and SUM44PE (SUM44)) and ER+ IDC (MCF7 and T47D). Unsupervised sample-to-sample correlation analysis segregated the ILC from IDC models, revealing a fundamental unique chromatin state of ILC (Figure 1A). Differential analysis (L2FC>1 and <−1, q-value<0.01) there were 11,777 sites significantly gained in ILC and 5,444 sites significantly gained in IDC (Figure 1B and S1A). The top enriched motifs in the ILC gained sites were FOXA1 motifs, followed by motifs of AP2*γ*, a key regulator of ER (16), and the estrogen response element (ERE) (Figures 1C and S1B). In contrast, the CTCF motif was the top motif in the IDC gained and the non-differentiated accessible sites (Figure 1D, S1C, S1D and S1E).

**Figure 1.**
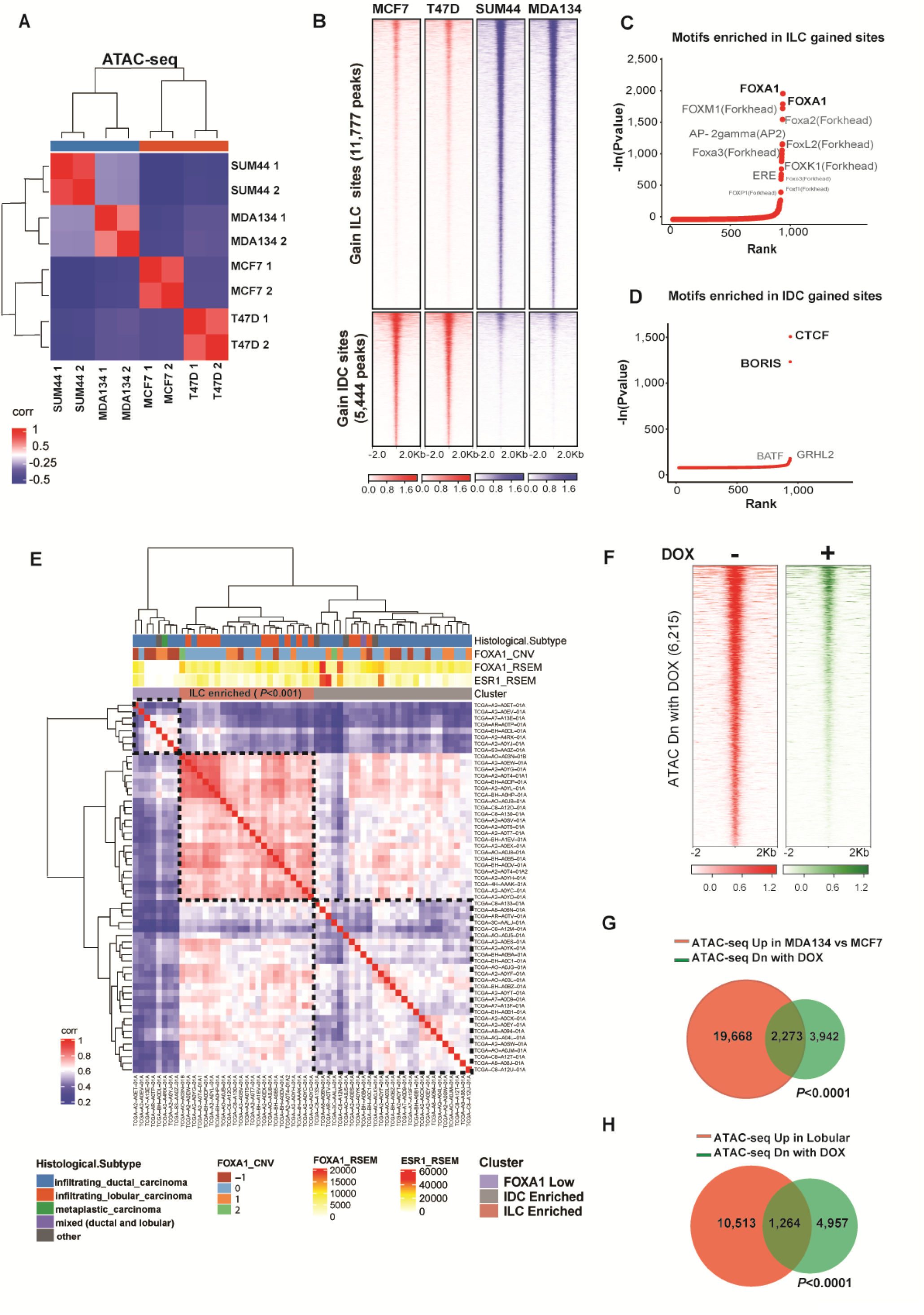
ILC has a unique chromatin cell state. **(A)** Sample to sample correlation of chromatin accessibility based on transposase-accessible chromatin followed by sequencing (ATAC-seq) by the Euclidean distance between rows/columns and Ward’s method of invasive lobular cancer (ILC) cells (MDAMB134 (MDA134) and SUM44) and invasive ductal cells (IDC) cells (MCF7 and T47D) cells after 10nM β-estradiol (E2) stimulation (cells were grown in hormone deprived (HD) conditions for 3 days followed by 45-minute treatment with 10nM E2). Shown in the plot are results of replicates. **(B)** Tornado plots of chromatin accessible sites gained in ILC cells (11,777 peaks) in blue and gained in the IDC cells (5,444 peaks) in red (Log 2FC >1 or <−1, *Q*<0.01). Chromatin accessible sites are shown in a horizontal window of ±2 kb from the peak center. **(C-D)** Ranking of motifs enriched in the ILC (**C**) and IDC (**D**) gained accessible sites based on p-value. **(E)** Sample to sample correlation heatmap of open chromatin sites in TCGA ER+ BC tumors applying only the chromatin accessible sites gained in the ILC cell line models (11,777 peaks). Samples are clustered by the Euclidean distance between rows/columns and Ward’s method. Samples cluster to three groups including an ILC enriched group (Fisher’s exact test). **(F)** Tornado plots of chromatin accessible sites lost when FOXA1 is downregulated by a doxycycline (DOX)-inducible shRNA after 3 days of DOX in presence of HD and 45min 10nM E2. **(G-H)** Venn diagrams of chromatin accessible sites upregulated in MDA134 in comparison to MCF7 (in red) **(G)** or upregulated in lobular cells in comparison to ductal cells (in red) **(H)** and the chromatin accessible sites lost by downregulation of FOXA1 by shRNA (in green).

To test the clinical relevance of these findings, we analyzed the ER+ BC samples of The Cancer Genome Atlas Program (TCGA) that were included in the Pan Cancer ATAC-seq study (17). This analysis consisted of 58 ER+ BCs with different histological subtypes (IDC: N=38, ILC: N=14 and other histological subtypes: N=6). None of these samples harbored FOXA1 mutations. Unsupervised sample-to sample correlation of the chromatin accessible sites gained in the ILC model cell lines segregated the samples to three main clusters: a FOXA1 low cluster (biologically an ER-negative cluster), ILC enriched cluster (including 11 of the 14 ILC samples, Fisher exact test p-value <0.001) and IDC enriched cluster (Figure 1E). Thus, ILC is characterized and can be identified by the unique chromatin accessible sites.

Because FOXA1 was the top motif enriched in the ILC gained chromatin accessible sites, we next tested the impact of FOXA1 silencing on chromatin accessibility by engineering MDAMB134 cells to stably express a doxycycline (DOX) inducible shFOXA1 (Figure S1F). Silencing of FOXA1 resulted in the loss of 6,215 accessible sites (L2FC <−1, q-value<0.01) without a gain in accessible sites (Figure1F). The lost sites significantly overlapped with the chromatin accessible sites gained in MDAMB134 compared to MCF7 cells or gained in ILC versus IDC cell models (Figures 1G and 1H).

Since we showed that FOXA1 has a role in facilitating the ILC unique chromatin state, we performed FOXA1 ChIP-seq in all four models and primary ILC cells obtained from a malignant peritoneal effusion from a patient with metastatic ER+ ILC. The majority of FOXA1 binding sites were in enhancer regions, including intronic and intergenic regions (Figure S2A). FOXA1 binding clustered the ILC models, including the primary ILC cells, and separated them from the IDC models (Figure 2A). This clustering was driven by 12,247 ILC gained peaks (L2FC>1, q-value <0.01) (Figure 2B). There was an insignificant number of peaks gained in IDC. As expected, FOXA1 motifs were the most significantly enriched motifs in the gained and non-differentiated peaks (Figure 2C, S2B and S2C). In addition, the ERE motif was among the enriched motifs (Figure 2). In support of the role of FOXA1 in the ILC unique chromatin state, the sites with gained chromatin accessibility in ILC had increased FOXA1 binding in ILC cells (Figure 2D). In addition, the chromatin accessible sites that were lost in the MDAMB134 cells after FOXA1 silencing had increased FOXA1 binding in MDAMB134 cells compared to MCF7 cells (Figure 2E). Strikingly, all the chromatin accessible sites gained in ILC overlapped with the ILC gained FOXA1 chromatin recruitment (Figure 2F). Moreover, there was a significant correlation between the intensity of the ILC gained FOXA1 binding and the intensity of gained chromatin accessibility in MDAMB134 and SUM44 cells (Spearman correlation coefficient=0.327, *P*<8.33e-98 for MDAMB134 and Spearman correlation coefficient=0.24, *P*<1.72e-50 for SUM44 (Figures 2G and S2D). In aggregate, ILC has a unique chromatin state that is interconnected with the reprogramming of FOXA1 recruitment.

**Figure 2.**
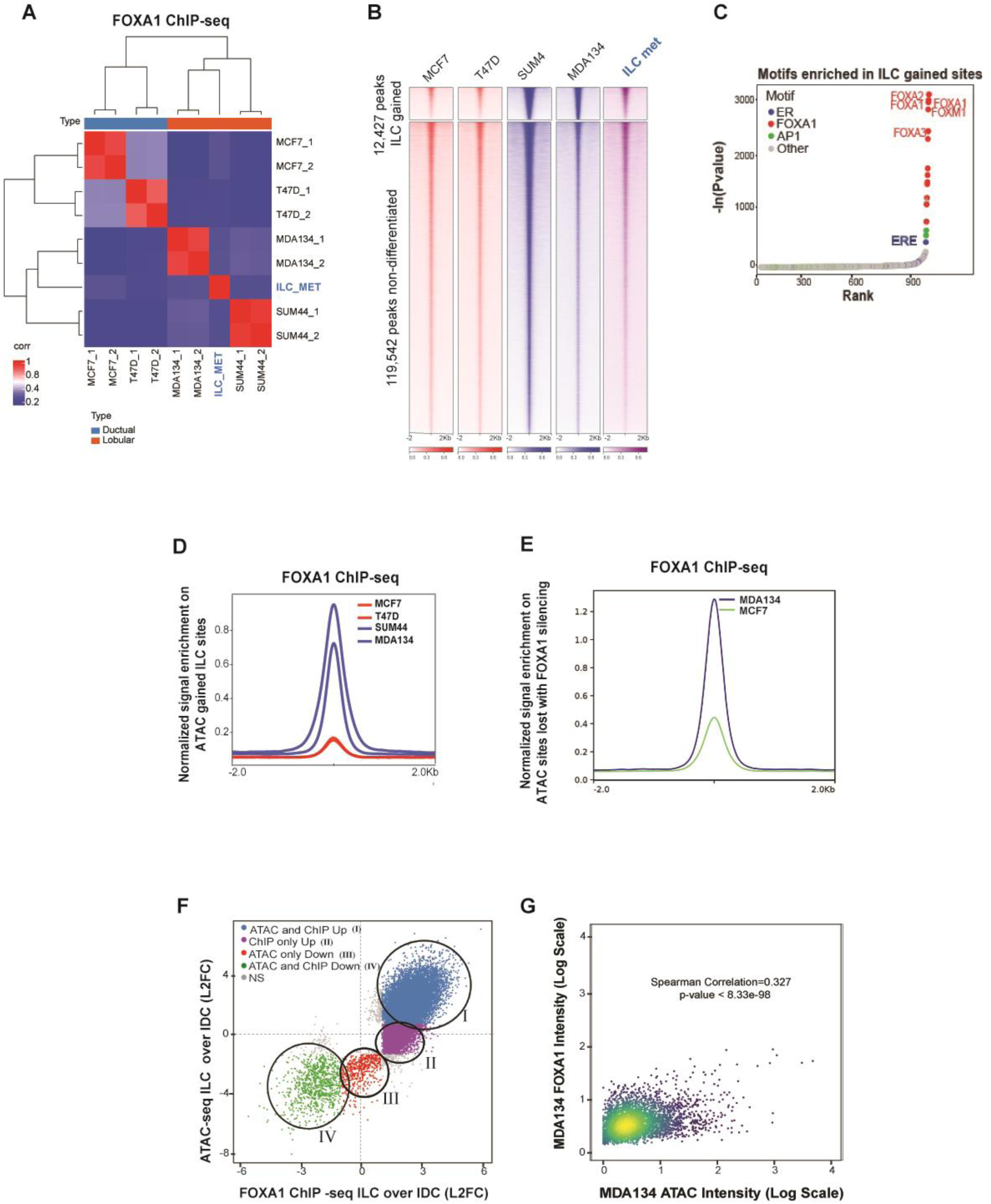
FOXA1 Reprogramming in ILC is linked to the ILC unique chromatin state. **(A)** Sample to sample correlation (Euclidean distance between rows/columns and Ward’s method) of FOXA1 binding sites correlation plots between all four cell lines in replicates (MCF7, T47D, MDA134 (MDAMB134) and SUM44) and the primary ILC cells isolated from a malignant peritoneal effusion from a patient with ER positive (ER+) metastatic ILC (ILC met). **(B)** Tornado plots of FOXA1 binding sites (12,427 sites) gained in ILC (MDA134 and SUM44) compared to IDC cells (MCF7 and T47D) and the union of the non-differentiated sites (Log 2FC >1 or <−1, *Q*<0.01). **(C)** Ranking of motifs enriched in the ILC gained FOXA1 binding sites based on p-value for enrichment. **(D)** Quantitative normalized signal of FOXA1 binding based on FOXA1 ChIP-seq in the ILC gained chromatin gained accessible sites based on the ATAC-seq analysis.**(E)** Quantitative normalized signal of FOXA1 binding based on FOXA1 ChIP-seq in the chromatin accessible sites lost in MDA134 after FOXA1 silencing based on the ATAC-seq analysis **(F)** Comparison of log2FC between ILC (MDA134 and SUM44) and IDC (MCF7 and T47D), FOXA1 binding sites (FOXA1 ChIP, x-axis) versus the log2FC comparing chromatin accessibility (ATAC-seq, y-axis) between ILC (MDA134 and SUM44) and IDC (MCF7 and T47D). **(G)** Intensity of binding in the intersecting sites of ILC gained FOXA1 binding and ATAC-seq in MDA134 versus MCF7 cells. Spearman correlation and p-value are reported.

### The ILC unique FOXA1 cistrome reprograms the ER transcriptional network

Given the known role of FOXA1 in facilitating ER binding(10–13), we next sought to determine how the ER axis is impacted by the reprogramming of the FOXA1 cistrome in ILC. To this end, we performed ER ChIP-seq in the ILC and IDC models. As shown previously (14), ligand dependent ER ChIP-seq of all four cell models showed that majority of ER binding sites were in enhancer regions (Fig. S3A). Unsupervised clustering of the ER binding sites segregated the ILC models, including the primary ILC cells isolated from a patient with metastatic peritoneal fluid, from the IDC models (Figure 3A). This clustering was driven by 6,885 peaks that were shared between MDAMB134 and SUM44 cells and gained compared to the IDC models (Log 2FC >1 or <−1, q-value <0.01) (Figure 3B). As expected, ILC ER gained binding sites were enriched in the ERE motif, followed by FOXA1 motifs (18) (Figures S3B and S3C). Importantly, comparison of ER with FOXA1 ChIP-seq showed that the ER binding sites gained in ILC had increased FOXA1 binding in ILC versus IDC, and there was a significant overlap between the ER sites gained in ILC and the FOXA1 ILC gained sites (44% of the gained ER binding sites overlapped with FOXA1 gained binding sites, p-value<0.0001) (Figure 3C and 3D). There were 9,367 ILC unique FOXA1 peaks that did not overlap with ER binding, which implies that FOXA1 has functions that are ER independent. Thus, the ILC gained FOXA1 binding sites are tightly linked to the ER gained binding but have functions that are independent of ER binding.

**Figure 3.**
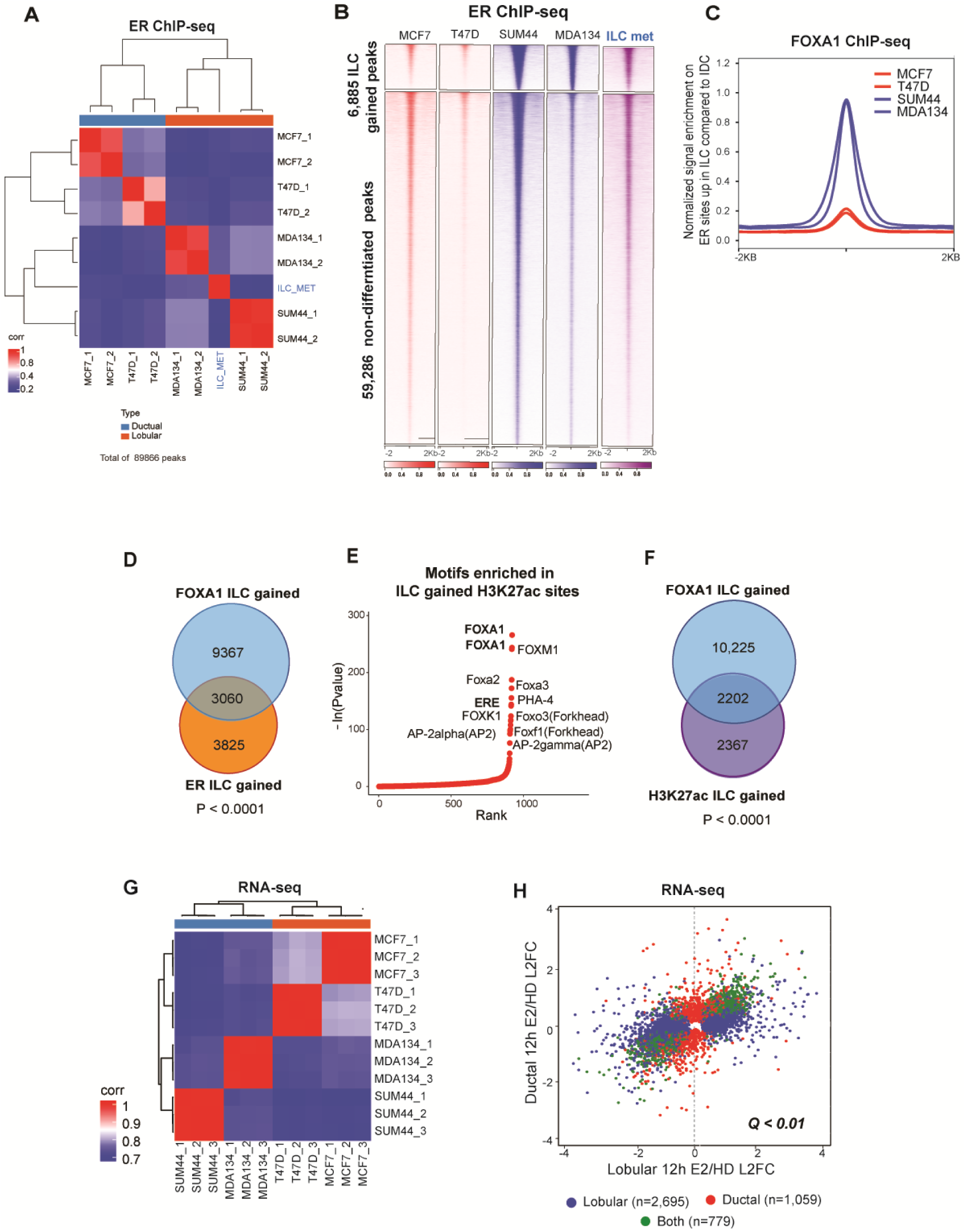
The ER cistrome in ILC cell lines models. **(A**) ER ChIP-seq sample to sample correlation plot. IDC (MCF7 and T47D), ILC (MDA134 and SUM44) and primary ILC metastatic cells isolated from malignant peritoneal effusion from a patient with ER+ metastatic ILC (ILC MET) samples are clustered by the Euclidean distance between rows/columns and Ward’s method. **(B)** Tornado plots of ER binding sites gained in ILC cells compared to IDC cells (6,885 peaks) and the union of the non-differentiated sites (59,286) (Log 2FC >1 or <−1, *Q*<0.01). ER binding sites is shown in a horizontal window of ±2 kb from the peak center. (**C)** Quantitative normalized signal of FOXA1 binding sites (FOXA1 ChIP-seq) on sites of ILC gained ER binding based on ER ChIP-seq in ILC cells (SUM44 and MDA134) and IDC cells (MCF7 and T47D). **(D)** Venn diagram of FOXA1 ILC gained peaks (12,427) in blue, and the ER ILC gained peaks (6,885) in orange, showing 3,060 peaks overlapping (sharing at least 1 bp) between the two factors. **(E)** Ranking of the motifs enriched in H3K27ac binding sites gained in ILC cells (SUM44 and MDA134) versus IDC cells (MCF7 and T47D) based on p-values of enrichment analysis (**F)** Overlap between FOXA1 ILC gained peaks (12,427) compared to IDC in blue, and the H3K27ac ILC gained peaks (4,569) compared to IDC in purple. **(G)** RNA-seq sample to sample correlation based on Euclidean distance between rows/columns and Ward’s method of IDC and ILC cells after hormone deprived (HD) and 12 hours of β-estradiol treatment. **(H)** Comparison of differentially expressed genes between hormone deprived (HD) and 12 hours of β-estradiol (E2) treatment in ILC (SUM44 and MDA134) and IDC (MCF7 and T47D) cells. Genes with differential expression of *Q*<0.01 are assigned to each category by the color scheme.

To start and elucidate the transcriptional significance of the ILC unique chromatin state and the reprogramming of the FOXA1-ER axis, we first performed H3K27ac ChIP-seq, a histone mark of active enhancers. Sample-to-sample correlation of H3K27ac ChIP-seq segregated ILC from IDC. This segregation was driven by 4,569 ILC gained sites (Figures S3D and S3E). The number of sites gained in IDC was limited. The top motif in the ILC gained H3K27ac sites was FOXA1 followed by ERE (Figures 3E, S3F and S3G)). Close to 50% of the ILC H3K27ac gained sites overlapped with the FOXA1 ILC gained sites (Figure 3F). Consistent with these findings, RNA-seq analysis of the four cell lines in estradiol stimulated conditions segregated the ILC from the IDC models (Figure 3G) and most of the estradiol regulated genes were different in ILC compared to IDC (Figure 3H). Taken together, these results support a previous study that showed disparate ligand stimulated transcription in ILC versus IDC (14) and provide initial evidence for the role of FOXA1 in the ILC unique transcription.

### The ILC-FOXA1 gene set is associated with increased risk of recurrence in ILC tumors of luminal A molecular subtype

To test if the ILC unique FOXA1 chromatin recruitment is driving the ligand stimulated transcriptional differences between ILC and IDC, we integrated the RNA-seq and ChIP-seq data applying Binding and Expression Target Analysis (BETA) (19). BETA basic showed that the FOXA1 binding sites gained in ILC are significantly associated with the upregulation of genes with increased expression in ILC versus IDC in estradiol stimulated conditions (Figure S4). This association is highlighted at the level of single genes in the volcano plot in figure 4a that shows the overlap between genes predicted to be gained by FOXA1 binding based on BETA minus and the genes upregulated in ILC versus IDC based on the RNAseq differential expression analysis (Figure 4A). To further substantiate the link between the FOXA1 gained sites and FOXA1 dependent transcription in ILC, we tested the association between the ILC gained binding sites and the genes regulated by the silencing of FOXA1 using the transcriptomic analysis of MDAMB134 cells with and without DOX induced silencing of FOXA1. BETA basic analysis showed that the ILC FOXA1 gained sites were significantly association with the genes that were down regulated after FOXA1 silencing (Figures S4C and S4D).

**Figure 4.**
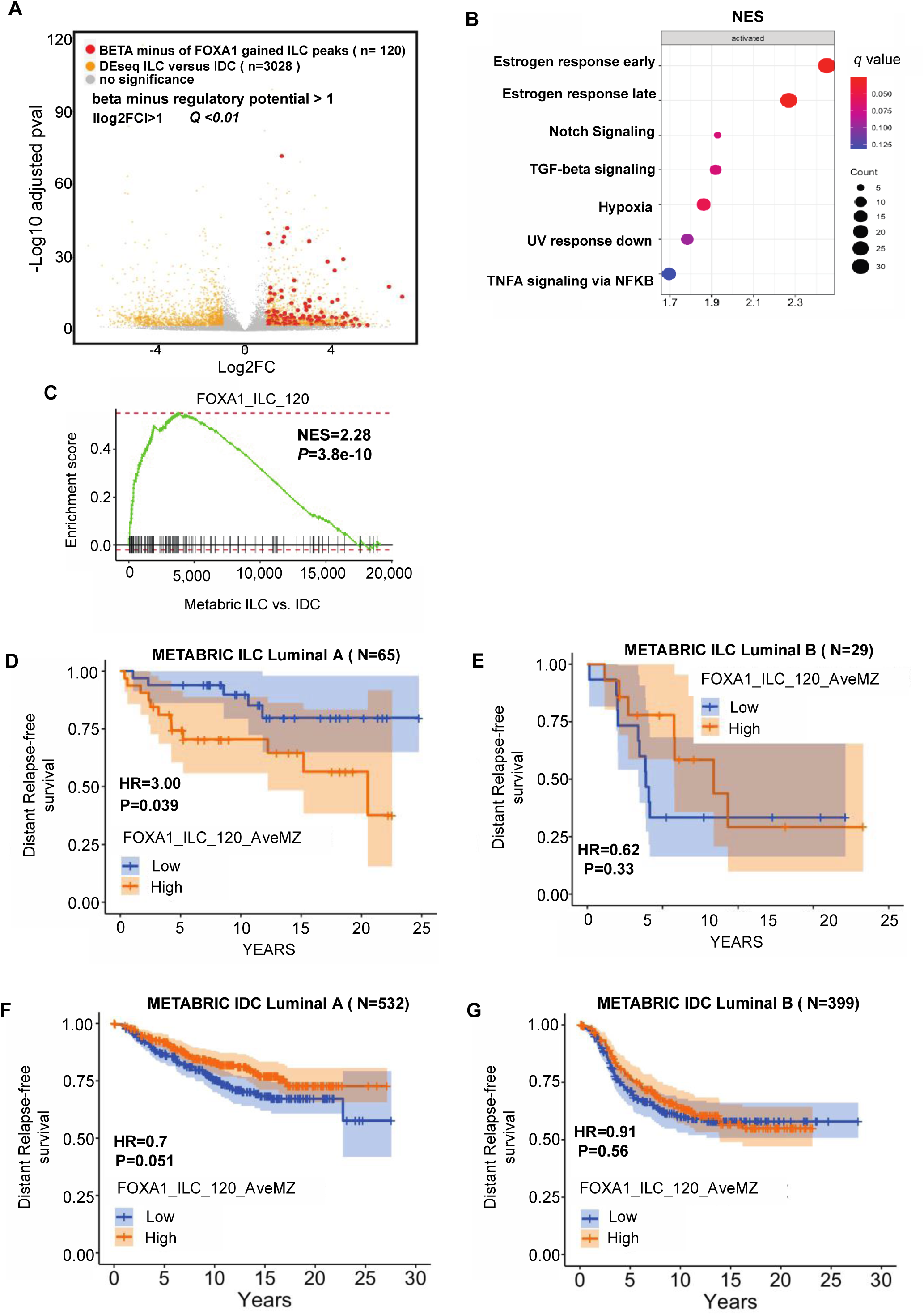
FOXA1 drives the ILC unique transcriptome. **(A)** Volcano plot depicting differential (DEseq2) expression comparing RNA-seq of ILC cells (MDA134 and SUM44) versus IDC cells (MCF7 and T47D) after β-estradiol stimulation (E2). Shown in yellow are the genes with significant differential expression ([log2FC] >1, *Q<*0.01. DESeq2). Red dots represent the genes with significant differential expression ([log2FC] >1, *Q<*0.01. DESeq2) and are regulated by FOXA1 ILC gained binding sites based on Binding and Expression Target (BETA) minus analysis. **(B)** Hallmark pathways enriched in the genes regulated by the ILC gained FOXA1 sites based on BETA basic. The normalized enrichment score (NES) is represented in the X-axis, the number of genes in the dataset is the count number represented by the circle size, *q*-value <0.25. (**C**) Enrichment plot from Gene Set Enrichment Analysis (GSEA) showing enrichment of the FOXA1_ILC_120 gene set derived from the ILC gained binding sites in ILC versus IDC estrogen receptor positive breast cancers in the METABRIC cohort. **(D)** Distant free survival in patients with Luminal A molecular subtype ILC from the METABRIC cohort comparing patients with high versus low expression of the FOXA1_ILC_120 gene set. **(E)** Distant free survival in patients with Luminal B molecular subtype type ILC from the METABRIC cohort comparing patients with high versus low expression of the FOXA1_ILC_120 gene set. **(F)** Distant free survival in patients with Luminal A molecular subtype IDC from the METABRIC cohort comparing patients with high versus low expression of the FOXA1_ILC_120 gene set. (**G**) Distant free survival in patients with Luminal B molecular subtype IDC from the METABRIC cohort comparing patients with high versus low expression of the FOXA1_ILC_120 gene set.

We next employed BETA basic to identify the genes that are direct targets of FOXA1 unique to ILC in estradiol stimulated conditions and generated a ranked product gene set termed the FOXA1-ILC gene set (Supplemental Table 1A). Ranked GSEA revealed that the FOXA1-ILC gene set is enriched in genes involved in key cancer pathways (ER response pathways, signaling (20,21), TGF-beta signaling (22,23), hypoxia (22), and NFKB signaling (24,25) (*Q*<0.25) (Figure 4B). We next generated a more stringent gene set of 120 FOXA1-ILC genes by selecting the genes with a ranked product p-value <0.001 (FOXA1_ILC 120). To study the clinical significance of this gene set we turned to the METABRIC cohort. We first determined that the FOXA1_ILC 120 is enriched in primary ER+ ILC tumors compared to primary ER+ IDC tumors (Figure 4C). Importantly, high versus low expression of the FOXA1_ILC 120 gene set was associated with a significantly lower distant relapse free survival (HR=3, p-value < 0.039) in patients with ILC and not IDC luminal A tumors. This association was also not seen in patients with luminal B breast cancers. Thus, the FOXA1_ILC 120 is a potential signature of high-risk ILC tumors that are driven by ER and not of tumors that are less dependent on ER signaling and are enriched in genetic alterations or other pathways associated with poor outcomes, such as P53 mutations (26) (Figures 4D,4E,4F and 4G). Hence, this signature could potentially identify the ILC patients that might benefit from improved endocrine treatment strategies.

We next looked at the overlapping FOXA1/ER/ H3K27ac sites gained in ILC and integrated these gained sites with the RNASeq differential expression comparing ILC and IDC in estradiol stimulated conditions by applying BETA basic to identify single genes that are direct targets of FOXA1 and ER and are upregulated in ILC. *CASZ1* was the top ranked gene, and *RET*, and *SNAI1* were among the top ranked genes that met these criteria (Supplemental Table 1B and Figure S4E). The protein levels of these three genes were increased in MDAMB134 versus MCF7 cells and downregulated by the silencing of ER or FOXA1 (Figure S4F). Previous studies have shown that RET and SNAIL are ER targets with roles in resistance to endocrine treatment (27–29). In contrast, CASZ1, a zinc finger TF that regulates transcription by binding to the nucleosome remodeling and histone deacetylase (NuRD) complex (30), has not been implicated as an important gene in BC. We first confirmed that CASZ1 expression is significantly higher in ER+ ILC versus ER+ IDC in the METABRIC cohort (Figure S4G). In the MDAMB134 model, silencing of CASZ1 resulted in cell growth inhibition (Figure S4H) and down regulation of genes with key roles in tumor growth, including *CDK2* and genes related to receptor tyrosine kinase signaling such as *FGFRL1, FGFR4, MAPK9* and *PIK3R2* (Figures S4I and S4J). Collectively, these results show that the ILC unique FOXA1 cistrome mediates the unique response to estradiol and the transcription of genes involved in tumor progression and poor outcomes.

### Down regulation of E-cadherin in IDC or up regulation of E-cadherin in ILC is not sufficient to induce a unique ER transcriptional program

Although, the loss of E-cadherin in ILC leads to the degradation of beta-catenin, p120 translocates to the cytoplasm or nucleus and in these compartments, it has a role in signaling and transcription regulation (31). We therefore asked if loss of E-cadherin and the release of membrane bound p120 contribute to the ILC unique ER cistrome and transcriptional response to estradiol. To test this, we silenced *CDH1* in MCF7 and T47D cells with a CRISPR cas9-gRNA (Figures S5A and S5B). The decrease in E-cadherin expression led to an increased migratory capacity but did not affect cell growth (Figures S5C and S5D). in addition, decreased E-cadherin expression resulted in limited transcriptional changes in MCF7 and T47D cells and the down regulated genes were involved in pathways of cell adhesion molecules among other pathways (Figures S5E,S5F,S5G and S5H). In addition, silencing of E-cadherin did not lead to significant changes in ER binding or H3K27 acetylation in estrogen stimulated conditions (Figures S5I and S5J). Consistent with these results, there were limited transcriptional changes after estrogen stimulation when comparing the cells with and without E-cadherin down regulation (Figures S5K and S5L). Specifically, FOXA1 and ER expression were not altered after E-cadherin silencing. Additionally, stable overexpression of E-cadherin in MDAMB134 cells with a DOX-inducible construct resulted in a limited number of transcriptional changes at 3 days and 3 weeks (Figures S5M, S5N and S5O). Taken together, silencing of E-cadherin by itself is not sufficient for the rewiring of the FOXA1-ER axis and altered estrogen response seen in ILC compared to IDC.

### A circuitry of FOXA1 upregulation mediated by a unique super-enhancer

Since we showed that aberrant FOXA1 activity is central to the epigenetic landscape in ILC, we sought to understand the mechanism of the altered FOXA1 recruitment in ILC. Mutations in TFs can contribute to altered binding and tumor progression (9,15), however, whole genome sequencing of the ILC models did not identify FOXA1 mutations (Figure S6A). The overexpression of TFs can also alter the regulatory activity of cancer cells (32) and, in fact, we observed mRNA and protein overexpression of FOXA1 in ILC compared to IDC (Figures 6A and S4F). Since we did not identify FOXA1 copy number variations or mutations in the promoter region to explain the upregulation of FOXA1 (Supplementary Fig. S6A) (21,33), we interrogated the TFs that regulate FOXA1 expression (cistromeDB toolkit (34)). Similar to other key lineage-defining TFs that have been found to be auto-induced (35), the TF with the highest FOXA1 regulatory potential was FOXA1 itself. ER and GATA3, the other two key determinants of ER+ BC, are also potent FOXA1 regulators (Figure 6B). Interestingly, examination of FOXA1 binding upstream to the FOXA1 transcription start site (TSS) revealed a FOXA1 peak in the enhancer region (>10Kb from the TSS) that was unique to the ILC models (P1) (Figure 6C). H3K27 acetylation in this region was also different in the ILC models compared to IDC and extended at least 3KB upstream in ILC overlapping with the P1 peak (Figure 6D). Based on the H3K27ac ChIP-seq analysis this region upstream to the FOXA1 promoter is a super-enhancer region in all four models (supplemental Table 2). However, the super-enhancer region in the ILC models extended beyond the IDC super-enhancer region and overlapped with P1 in the ILC models only. We also detected the ILC unique FOXA1 P1 peak and super-enhancer region in ILC metastatic cells isolated from malignant peritoneal fluid, demonstrating clinical relevance of the P1 peak and unique super-enhancer region (Figures 6C and 6D). Since super-enhancers are characterized by the binding of multiple tissue-specific TFs (36) we looked at the binding of ER and GATA3, the other top ranked FOXA1 regulators, and saw increased ER and GATA3 binding in the ILC unique super-enhancer region (Figure 6E). Furthermore, H3K27ac Hi-ChIP detected ILC unique loops in this region (Figure S6B). Next, we tested the regulatory function of P1 by selective targeting of P1 with dCAS9-KRAB and a gRNA targeting the summit of this peak (details of the coordinates of this site are in the methods). In the SUM44 ILC cells inactivation of P1 decreased the expression of FOXA1 and led to the regression of cell growth (Figures 5F, 5G and 5H). In contrast, inactivation of P1 in MCF7 cells did not impact FOXA1 expression or cell growth despite the sensitivity of MCF7 to a siFOXA1 (Figures 5I, 5J and S6C). The activity of dCAS9-KRAB in MCF7 cells was confirmed by infecting the MCF7 cells engineered to stably express dCAS9-KRAB with a BFP-GFP reporter construct that includes a gRNA that targets the GFP cassette. In these cells, expression of dCAS9-KRAB along with the gGFP resulted in the loss of expression of BFP and GFP (Fig. S6D). Collectively, our results support FOXA1 overexpression in the ILC models through auto-induction by a unique FOXA1 binding site within a super-enhancer.

**Figure 5.**
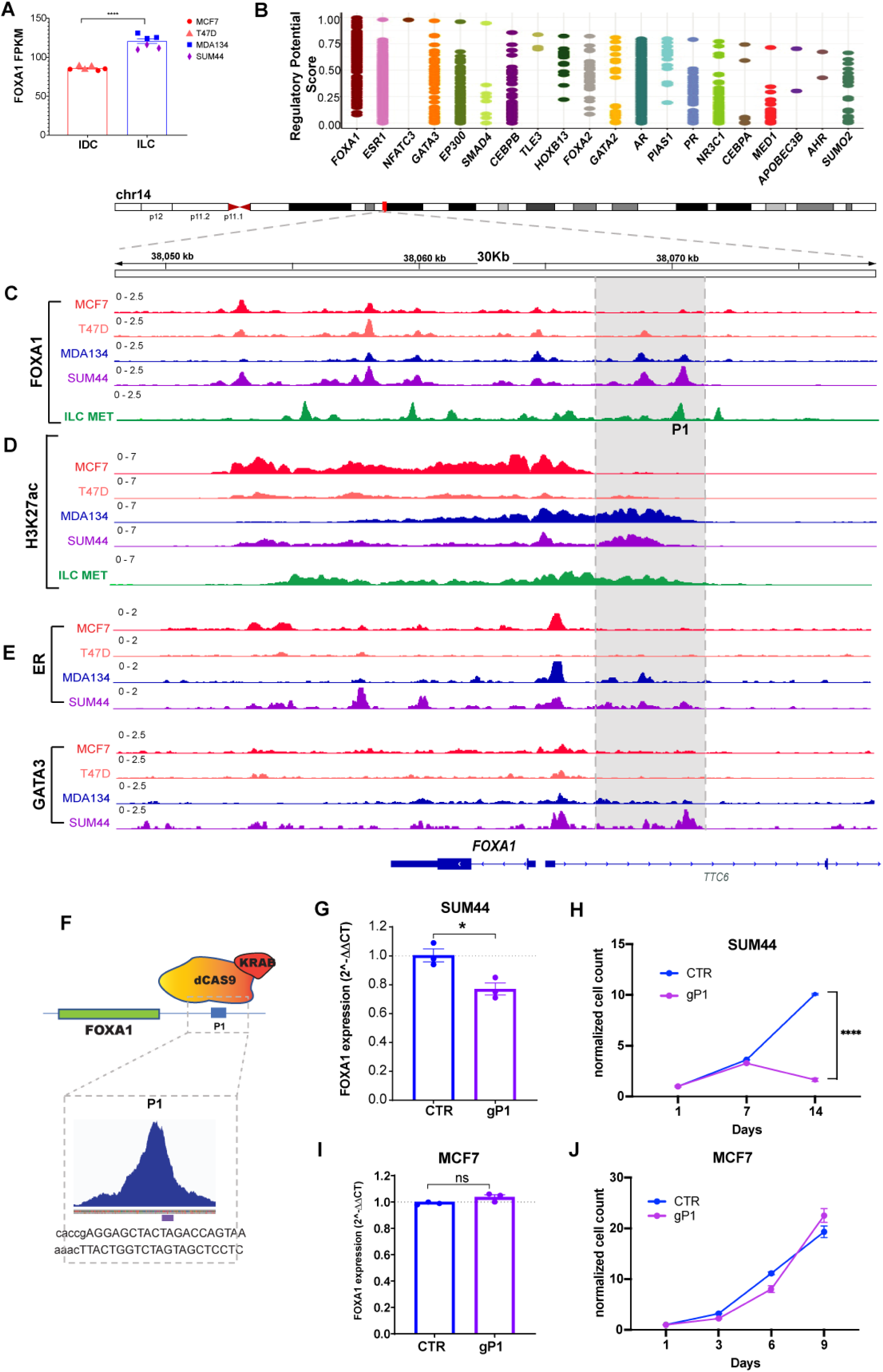
FOXA1 auto-induction in ILC through a unique super-enhancer and FOXA1 binding site. **(A)** RNA expression (FPKM values) of FOXA1 in IDC (MCF7 and T47D) and ILC (MDA134 and SUM44) cells. ****, t-test p-value <0.00001 **(B)** Regulatory potential of the transcription factors that regulate FOXA1 based on the CistromeDB toolkit (dbtoolkit.cistrome.org). **(C-D)** ChIP-seq tracks showing FOXA1 **(C)** and H3K27acetylation **(D)** in the IDC (MCF7 and T47D (red and orange tracks) and ILC (MDA134 and SUM44) (blue and purple tracks) cell lines and primary cells from ILC metastasic peritoneal effusion (ILC met, green tracks). **(E)** ER and GATA3 ChIP-seq tracks for the IDC (MCF7 and T47D) (red and orange tracks) and ILC (MDA134 and SUM44) (blue and purple tracks) cell lines. **(F)** Cartoon of CRISPRi action at the P1 site. Enlargement showing the gRNA used to target the FOXA1 binding region. **(E)** mRNA levels of FOXA1 in SUM44 lobular cells by reverse transcription quantitative PCR (RT qPCR) in presence of the control guide and the guide RNAs targeting P1 (gP1). The mRNA expression was normalized to the GAPDH housekeeping gene, and expression levels are presented as 2^–ΔΔCT compared with control. **(F)** Cell proliferation assay after 14 days in SUM44 cells of control cells and after suppression of the FOXA1 P1 (gP1) using CRISPRi. Error bars represent ±SEM, n=3. **(G)** FOXA1 mRNA levels of MCF7 in presence of the control guide and the guide RNAs targeting P1 (gP1). **(H)** Cell proliferation assay in MCF7 cells in presence of the control guide and the guide RNAs targeting P1 (gP1). *p < 0.05; ****p < 0.0001; ns: not significant.

**Figure 6.**
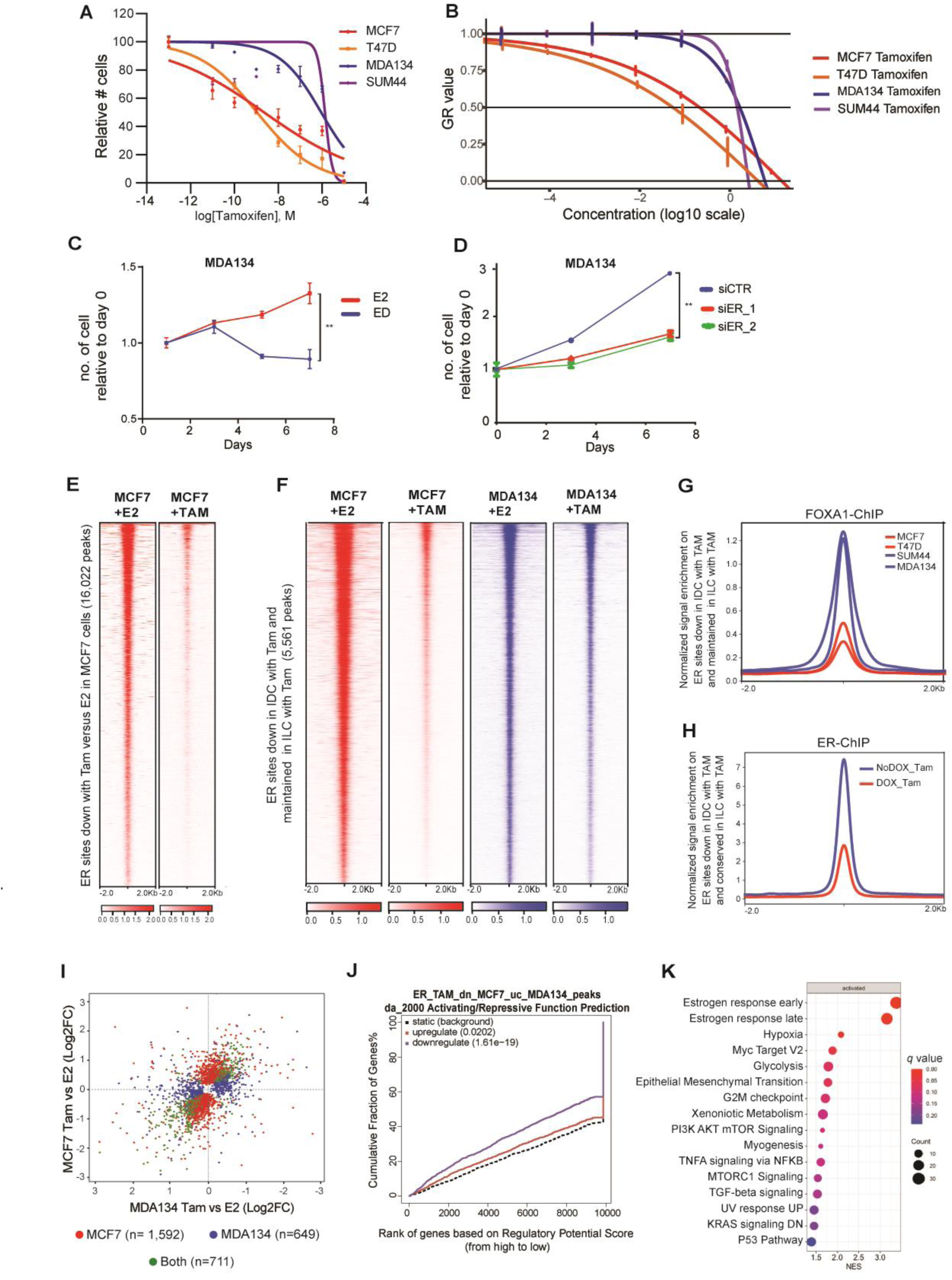
The mechanism of Tamoxifen Resistance in ILC. **(A-B)** Dose response curves of 4-hydroxytamoxifen (tamoxifen) treatment in ductal and lobular cell lines. (**A**) Curves normalized to vehicle control. IC50 values is in the range of 1uM for both ILC models (MDAMB134 (MDA134) and SUM44), compared to an IC50 of 1nM and 2.2nM for T47D and MCF7, respectively. **(B)** GR50s values are 240nM for MCF7 60nM for T47D, 1920nM for MDA134 and 1680µM for SUM44. **(C)** Cell proliferation curves of MDA134 cells followed for seven days without (ED) or with estadiol (E2, 10nM). Error bars represent ±SEM, n = 3. **, p value <0.01 **(D)** Cell proliferation studies of MDA134 cells in full medium conditions including cells transfected with an siControl (siCTR) and cells with silencing of ER (siER_1 and siER_2). Error bars represent ±SEM, n = 3. ****, p value <0.0001 **(E)** Tornado plots of ER binding sites lost in MCF7 treated with 4-hydroxytamoxifen (TAM) compared to β-estradiol (E2) treatment. (**F)** Tornado plots of ER binding sites lost in MCF7 treated with TAM compared to E2 treatment and unchanged in the MDA134 cells in E2 and TAM treated conditions (Log 2FC >1 or <−1, q-value<0.01). **(G)** Quantitative normalized signal of FOXA1 ChIP-seq binding at ER binding sites lost in MCF7 cells with TAM treatment but unchanged in MDA134 cells. **(H)** Quantitative normalized signal of ER ChIP-seq binding on ER sites lost in MCF7 cells but retained in MDA134 cells, in MDA134 cells treated with TAM (with and without DOX induction of FOXA1 silencing (shFOXA1). **(J)** Comparison of differentially expressed genes between β-estradiol (E2) and 4-hydroxytamoxifen (Tam) in MCF7 and MDA134. Genes with FDR <0.05 are assigned to each category by the color scheme. There were 711 shared differentially expressed genes, 1,592 genes differentially expressed in MCF7 cells only, and 649 genes differentially expressed exclusively in MDA134. **(K)** Binding and Expression Target Analysis (BETA) basic plot of the activating and repressive function of the ER binding sites lost in MCF7 but conserved in MDA134 after tamoxifen treatment. The red line represents the genes upregulated and the purple line the genes downregulated with 4hydroxytamoxifen versus β-estradiol treatment in MCF7 cells. The black dashed line indicates the non-differentially expressed genes as background. The p-value is based on the Kolmogorov-Smirnov test. **(L)** Hallmark pathways enriched in the genes determined to be downregulated by 4-hydroxytamoxifen compared to E2 treatment and regulated by the ER binding sites lost in MCF7 cells after tamoxifen treatment but unchanged in MDA134. The normalized enrichment score (NES) is represented in the X-axis, the number of genes in the dataset is the count number represented by the circle size, *q*-value <0.25.

### The reprogrammed FOXA1 cistrome drives Tamoxifen resistance in ILC

Similar to the clinical observation of relative resistance to tamoxifen in ILC (6), MDAMB134 and SUM44 cells were resistant to tamoxifen (IC50 of close to 1uM for both ILC models compared to an IC50 of 1nM and 2.2nM for T47D and MCF7, respectively) (Figure 6A). Since the ILC models have a longer doubling time compared to the IDC models, we confirmed the resistance to tamoxifen by growth rate metrics, which are independent of the division rate of the assayed cells (37). The GR_50_ for tamoxifen was 8-30-fold higher in the ILC models compared to IDC (Figure 6B). Tamoxifen resistance was not due to ligand or ER independent growth, as evidenced by the high sensitivity of MDAMB134 cells to estrogen deprivation and ER silencing, as in MCF7 cells (Figures 6C, 6D, S7A and S7B).

To start to understand the mechanism of relative resistance to tamoxifen in ILC, we performed ER ChIP-seq after tamoxifen or estradiol treatment in MCF7 and MDAMB134 cells. In MCF7 cells, tamoxifen compared to estradiol led to the loss of 16,022 ER binding sites, whereas in MDAMB134 cells there were 37 lost and 236 gained ER binding sites (Log 2FC >1 or <−1 and q-value<0.01) (Figures 6E and S7C). When comparing the two cell lines and two treatment conditions, we identified 5,561 peaks that were lost in MCF7 cells but retained in MDAMB134 cells after tamoxifen treatment (Log 2FC >1 or <−1 and q-value<0.01) (Figure 6F). These ER peaks had increased FOXA1 binding and chromatin accessibility in ILC cells compared to IDC cells (Figures 6G and S7D). Furthermore, down regulation of FOXA1 expression with DOX-induced shFOXA1 in MDAMB134 cells (Figure S1F) resulted in growth inhibition and a loss of ER binding in the sites maintained after tamoxifen treatment (Figures S7E and 6H). Of note, we showed that FOXA1 silencing leads to decreased ER expression (S4F), hence, these functional consequences of FOXA1 silencing can be attributed to its role in facilitating ER binding and regulating the expression of ER.

To study the transcriptional consequences of the disparate effects of tamoxifen treatment on ER binding in ILC versus IDC, we performed RNA-seq with and without tamoxifen treatment in MCF7 and MDAMB134 cells. In line with the differences in the changes in ER binding in response to tamoxifen treatment, there were differences in the transcriptional changes in response to tamoxifen treatment. Comparison of the transcriptional changes induced by tamoxifen in MCF7 and MDAMB134 revealed 1,592 genes differentially expressed in MCF7 cells only, 649 genes differentially expressed in MDAMB134 only, and 711 genes differentially expressed in both cell models (Figure 6I). BETA basic analysis showed that the ER binding sites lost in MCF7 cells but retained in MDAMB134 after tamoxifen treatment had a significant association with the genes downregulated with tamoxifen treatment in MCF7 cells (Figure 6J). GSEA of the ranked genes regulated by the MDAMB134 tamoxifen-maintained ER binding sites in MCF7 cells showed that these genes are involved in pathways that are key therapeutic targets of tamoxifen, such as response to estrogen, MYC targets, and epithelial mesenchymal transition (38) (Figure 6K, supplemental table 1C). In aggregate, in MCF7 cells tamoxifen treatment results in the loss of a set of ER binding sites and this is associated with the downregulation of genes important for the therapeutic response to tamoxifen. In contrast, in MDAMB134 cells, ER binding at these sites is retained in a FOXA1 dependent manner.

## Discussion

In this study, we found that ILC has a unique chromatin state associated with the altered of FOXA1 recruitment. The ILC unique epigenetic state provides one mechanism to explain the disparate transcriptional responses to estrogen and resistance to tamoxifen despite the high ER expression and ER dependency. These results provide new fundamental insights to the unique biology of ILC and have important therapeutic implications.

Several mechanisms of aberrant FOXA1 signaling have been described previously, including FOXA1 mutations and amplifications (7-9,21). Though the increased frequency of FOXA1 mutations in ILC suggests that altered FOXA1 signaling is important in the biology of ILC, the majority of ILCs do not harbor a FOXA1 mutation. This implies that other mechanisms of aberrant FOXA1 signaling may be enriched in ILC. Indeed, we identified another mechanism of altered FOXA1 signaling in cell lines and metastatic ILC cells isolated from a patient with a metastatic peritoneal effusion. Herein, we provided evidence for a feed-forward circuit of FOXA1 auto-induction involving a unique super-enhancer and FOXA1 binding site with the co-recruitment of other TFs including ER and GATA3. This circuit demonstrates how the overexpression of a single key TF can lead to sustained effects on an entire transcriptional program and facilitates tumor progression and treatment resistance.

Previous studies identified several mechanisms of tamoxifen resistance unique to ILC models (39,40). We have now unraveled a novel mechanism of tamoxifen resistance in ILC. Tamoxifen is a selective estrogen receptor modulator that inhibits the transcriptional activity of ER by shifting the balance between co-activator and co-repressor binding (41). Resistance specific to tamoxifen has been attributed mainly to a shift towards increased co-activator versus co-repressor binding, which can lead to ER agonistic activity and explain tumor regression after tamoxifen withdrawal (42,43). Here we show that in MCF7 cells tamoxifen treatment leads to loss of ER binding in selective sites and reduces ER mediated transcription. Our results suggest that gained FOXA1 recruitment in ILC can counteract this activity of tamoxifen and preserve ER binding. In addition, FOXA1 silencing was sufficient to inhibit this ER binding either through downregulation of ER expression or by the perturbation of ER binding. Together, these results highlight the importance of developing approaches for targeting traditionally undruggable targets such as FOXA1.

We provide proof of principle for potent tumor growth inhibition by selective targeting of a unique FOXA1 binding site associated with a super-enhancer that regulates the expression of FOXA1 itself. The advantage of this type of therapeutic approach over current epigenetic strategies is twofold: 1. Substantial efficacy because of the disruption of an auto-regulatory loop that is self-sustained and regulates multiple genes involved in tumor progression. 2. High precision given the disruption of a selective super-enhancer in selective cells as opposed to current therapeutic approaches that target epigenetic regulators and modulate the transcription of multiple genes in a non-selective manner (44). Importantly, selective inhibition of a regulatory region with a small molecule has been shown to be feasible (45). However, of more immediate clinical relevance, we show that ILC is highly dependent on a unique ER transcriptional axis and cell growth remains ligand dependent. These results support pre-clinical studies of ILC models and clinical trials dedicated to patients with ILC to investigate the oral selective estrogen receptor degraders and other novel endocrine treatments currently in clinical development.

## Acknowledgements

We would like to Ms. Cheryl Fox for her support, important insights and helpful discussion.

## Author contributions

Conceptualization: A.N., O.M.F., E.P.W, H.L., M.L.F., R.J., R.S.

Data curation: A.N., S.S., G.C.F., M.P, W.L., A.F-T., C.G, F.H-P., S.S., K.L., H.M, A.O.

Formal Analysis: X.Q., A.F., A.F., Y.X, M.C, A.N, X.F, R.S, J.B.

Funding acquisition: R.J, M.L.F., E.P.W

Investigation: A.N., S.S., G.C.F., M.P, W.L., A.F-T., C.G, F.H-P., S.S., K.L., H.M, A.O.

Methodology: A.N., R.J., H.L.,M.F.

Resources: E.P.W.

Software: X.Q., A.F., A.F., Y.X, M.C.

Supervision: E.P.W, R.J, M.L.F., H.L., O.M.F.

Validation: A.N., R.J.

Visualization: A.N., A.F., A.F.

Writing – original draft: R.J., A.N.

Writing – review & editing: All authors

