## Supplemental methods and figures for "A Distinct Chromatin State Drives Therapeutic Resistance in Invasive Lobular Breast Cancer"

Supplemental References

Supplemental Figures 7

### Supplemental Methods

#### Proliferation assays

The breast cancer cells were plated in 24 well plates. At indicated time points, the cells were counted using the Celigo image Cytometer (Nexcelom). Hoechst was used for nuclear staining and propidium iodide was used for staining dead cells. 4-hydroxytamoxifen (Sigma-Aldrich) was used for the dose response studies and determination of the IC50's and GR50's (1) at 7 days.

#### Migration assay

CytoSelectTM 24-Well Cell Migration (Cell Biolabs, INC) was used as per manufacturer instruction. Briefly, 300,000 cells were placed in the upper chamber in serum free media and incubated for 48 hours. Migratory cells moving towards the lower chamber containing medium with 10% FBS were stained after removing the non-migratory cells in the upper chamber. Pictures of the stained cells were used to be analyzed and quantified with ImageJ software.

#### Western blotting

Cells were lysed in RIPA buffer (Boston BioProducts) supplemented with protease and phosphatase inhibitors and subjected to SDS-PAGE. Antibodies used include: E-Cadherin (24E10), Ret (E1N8X), Snail (C15D3), and CDK2 (78B2), from Cell Signaling Technology, CASZ1 (600-401-B62) from VWR, FOXA1 (ab23738) from Abcam, ERα (sc-543) and GAPDH from Santa Cruz. Western blotting bands intensity were quantified by ImageJ.

RET: https://antibodyregistry.org/search?q=RRID:AB_2798509

SNAIL: https://antibodyregistry.org/search?q=RRID:AB_2255011

CDK2: https://antibodyregistry.org/search.php?q=AB_2276129

CASZ1: https://antibodyregistry.org/search.php?q=AB_1961495

FOXA1: https://antibodyregistry.org/search.php?q=AB_2104842

FOXA1: https://antibodyregistry.org/search.php?q=AB_304744

ER: https://antibodyregistry.org/search.php?q=AB_631471

ER: https://antibodyregistry.org/search.php?q=AB_61659

GAPDH: https://antibodyregistry.org/search.php?q=AB_10167668

**Immunofluorescence**

Cells were cultured on glass slides for 48hrs. Cells were fixed and permeabilized 10 min at room temperature (RT) with Methanol-Acetone 1:1, blocked in 2% bovine serum albumin (BSA) in PBS + 0.1% Triton X-100 (PBST) for 30 minutes, and then incubated with anti-CDH1 (24E10), anti-β-catenin (D10A8), or anti-p120-catenin (clone 98) primary antibody diluted in 2% BSA in PBS, overnight at 4°C. After four washes in PBST, samples were incubated with secondary antibodies (AlexaFluor-488 or AlexaFluor-594, 1:300 in 2% BSA in PBST) for 1h RT in darkness. Nuclei were counterstained with DAPI (Life Technologies). Images were acquired with a fluorescence microscope equipped with a color camera.

**Generation of knock-out cells (CRISPR, ShRNA and siRNA)**

**-CRISPR/Cas9 cells**

Construction of lenti-CRISPR/Cas9 vectors targeting E-cadherin (CDH1) in the two ductal cell lines was performed following the protocol associated with the backbone vector (49535, Addgene). The following sgRNA for CDH1 sequences were used:

gRNA1 F:5′ CACCGAGCTGGCTGACATGTACGG 3′

gRNA1 R: 5′ AAACCCGTACATGTCAGCCAGCTC 3′

**- DOX inducible ShFOXA1**

FOXA1 shRNA constructs (provided by Novartis) were generated by inserting annealed oligonucleotides into EcoRI/AgeI-digested pLKO-Tet-On plasmid.

The FOXA1 sequence to target was: 5' GCAGATGTCTTTAAATGAAAT.

Top oligo:

CCGGGCAGATGTCTTTAAATGAAATCTCGAGATTTCATTTAAAGACATCTGCTTTTT

Bottom oligo:

AATTAAAAGCAGATGTCTTTAAATGAAATCTCGAGATTTCATTTAAAGACATCTGC

Lentivirus was produced in 293T cells to infect cells in media containing polybrene (8-10µg/mL). Cells were selected with puromycin treatment after transduction. 1ug/mL of doxycycline was used for the induction of the shFOXA1.

**-siRNA**

MCF7 and MDA134 cells were seeded in 6- or 24-well plates and transfected with 20nM siRNA oligos by Lipofectamine RNAiMax reagent (Life Technology). Knockdown efficiency was determined after 72 hours of transfection. Cell counting was performed after 3 and 7 days of transfection. The ON-TARGET siRNA oligos targeting ESR1 (LQ-003401-00), FOXA1 (J-010319-05 and J-010319-06), CASZ1 (LQ-020764-02) or non-targeting control (D-001810-01) were all purchased from Dharmacon.

**RNA isolation and reverse transcriptase quantitative PCR analysis**

Total RNA was isolated using TRIzol (Life Technologies) and RNeasy Mini Kit (Qiagen), and reverse transcribed as previously described. Quantitative real-time PCR amplification was performed using Lightcycler 480 SYBR Green I Master (Roche Diagnostics) as per manufacture's instruction and primers for FOXA1 and GAPDH were:

FOXA1_For GAAGATGGAAGGGCATGAAA

FOXA1_Rev GCCTGAGTTCATGTTGCTGA

GAPDH_For CGAGATCCCTCCAAAATCAA

GAPDH_Rev TTCACACCCATGACGAACAT

The relative fold differences in gene expression were calculated by the $2^{-\Delta\Delta Ct}$ method over the control and with GAPDH as a normalization control.

**ATAC-seq**

For the global chromatin accessibility experiments ductal and lobular cell lines were cultured for three days in HD conditions and then treated with 10nM estradiol for 45 minutes. Nuclei of 100,000 freshly fixed in 1% formaldehyde cells were processed as previously reported (2). Briefly, cells were resuspended in 1 mL of cold ATAC-seq resuspension buffer (RSB; 10 mM Tris-HCl pH 7.4, 10 mM NaCl, and 3 mM MgCl2 in water) and centrifuged. Cell pellets were then resuspended in ATAC-seq RSB (0.1% NP40, 0.1% Tween-20, and 0.01% digitonin) and incubated on ice. After lysis, ATAC-seq RSB containing 0.1% Tween-20 (without NP40 or digitonin) was added. Nuclei were centrifuged and then were resuspended in 50 µl of transposition mix (25 µl 2× TD buffer, 2.5 µl transposase~~26~~ (100 nM final), 16.5 µl PBS, 0.5 µl 1% digitonin, 0.5 µl 10% Tween-20, and 5 µl water). Transposition reactions were incubated at 37 °C for 30 min. Reactions were cleaned up with Zymo DNA Clean and Concentrator 5 columns.

**Hi-ChIP**

Hi-ChIP libraries were generated as per the method of Mumbach et al.(3) with modifications. In brief, nuclear isolation and in situ Hi-C contact generation were performed by proximity ligation on 5-10 million cross-linked cells per condition, using biotin-dATP. Next, nuclei were lysed and sonicated. 20ug of chromatin were used for immunoprecipitation against the epitope of interest (H3K27AC C15410196, Diagenode). Next, reverse cross-linking, DNA purification and biotin capture of the Hi-C contacts were performed prior to DNA library preparation with Tn5 on beads, amplifying for 11 cycles total. Libraries were sequenced on an Illumina platform.

Hi-ChIP reads were aligned to the human reference genome (hg19) using bowtie2 version 2.3.4.3 and preprocessed in HiC-Pro version 2.9.0(4). Default settings were used to remove duplicate reads, assign reads to DpnII restriction fragments and filter for valid interactions. The fit HiChIP pipeline version 7.1(5) was used to process all valid reads from HiC-Pro. For significant contact calling, all read pairs must have at least one anchor in the H3K27ac ChIP-seq peaks. Chromatin interactions were filtered using a minimum distance of 5 kb and a maximum of 3 Mb. The final set of chromatin loops used for further investigation were interactions which were selected by q-value <1%.

**ChIP sequencing analysis**

ChIP reads were aligned to the hg19 genome assembly using BWA (6) and ChIP-seq peaks were called using MACS 2.0 (7,8). For differential analysis, bed-files were merged using bedops (9) and then DEseq2 was used to assign differential intensities and statistics. Differential binding sites and open chromatin was determined by filtering out insignificant peaks (q-value <0.01) and then determining log2FC values between samples for each region using a log2 FC >1 or <-1, q-value<0.01. Unsupervised sample-to-sample correlation analysis was done using Euclidean distance and Ward's method. Analysis was done using the CoBRA workflow (10). The ILC or IDC unique binding sites and the union of the non-differential binding sites are plotted in the Tornado plots.

For integration of RNA-seq and ChIP-seq, we used the Binding and Expression Target Analysis (BETA) (11). This tool integrates ChIPseq and RNASeq to infer direct transcriptional targets and the significance of transcriptional function of a specific transcription factor. We used 2 components of this tool, including: 1. BETA basic (transcription factor target gene prediction based on a regulatory score applying ChIPseq data only), 2. BETA basic (BETA) includes: (a) transcription factor activating, or repressive function prediction based on ChIPseq and RNAseq data using the Kolmogorov-Smirnov test (b) a ranked product list of genes (ranking based on the

ChIP data and differential expression data) with a p-value for each gene (the probability that the gene is regulated by a specific transcription factor and is differentially expressed).

The significance of overlapping sites was tested using Fisher's exact test in GeneOverlap (1.30.0) R package. The motif analysis was done using HOMER (v4.4) (12). The Rank Ordering of Super Enhancers was used (ROSE) was applied for identification of super enhancers (13,14).

**TCGA ATAC-seq**

The breast cancer TCGA ATAC-seq data was downloaded from the Genomic Data Commons Data Portal (dbGAP study accession phs000178). ER + breast cancer samples were analyzed as outlined on the ATAC-seq- analysis section in the main methods section. Samples were clustered by the Euclidean distance between columns and Ward's method.

**RNA sequencing analysis**

The RNA-seq analyses were performed using the VIPER analysis pipeline (15,16). Alignment to the hg19 human genome was done using STAR v2.7.0f followed by Transcript assembly using cufflinks v2.2.1 (17) and RseQC v2.6.2 (18). Differential Expression Analysis was done using DEseq2 v1.18.1 (19). GSEA analysis was performed using the Broad GSEA Application (20).

**Survival Analysis**

**Calculation of Average Modified Z-score (AveMZ)**

Breast tumor gene expression profiles with clinicopathologic data were obtained from METABRIC (21,22) via authorized access in Synapse (synapse.sagebase.org). To calculate the

FOXA1-ILC_120 gene set expression in the METABRIC data set we used the modified Z-score to transform the gene expression data because it relies on the median and is less influenced by outliers when compared to the standard Z-score based on the mean. The score was calculated by the formula of ($X$-MED)/(1.486*MAD), where $X$ is the log-transformed gene expression value. MED is the median level of $X$ across samples, and MAD ($\neq 0$) is the median absolute deviation calculated by MAD = median ($|X$ - median$(X)|$). The signature score of target gene set was presented as AveMZ by calculating the mean of modified Z-scores of the signature genes.

Kaplan-Meier plots were produced using the R survminer v0.4.8 package to display the survival probabilities per group as a function of time. Groups of ER+ BC cohorts were stratified by the median cutoff of the gene signature score. Hazard ratio (HR) of stratum and $P$ value were calculated using the Wald estimates and likelihood ratio test, respectively, in the Cox proportional hazards model using the R survival v3.2-3 package.

### Supplemental Figures

**Figure S1. ATAC-seq in invasive ductal cancer and invasive lobular cancer models.**

**(A)** Comparison of open chromatin sites in invasive lobular cancer (ILC) and invasive ductal cancer (IDC). The plot depicts the sites gained in the ILC cells (11,777 peaks), gained in the (IDC) cells (5,444 peaks), and the union of the sites not differentiated between IDC and ILC cells (96,508 peaks). Open chromatin sites are shown in a horizontal window of ±2 kb from the peak center. **(B)** Table of the top enriched motifs in the chromatin accessible sites gained in ILC. **(C)** Ranking of motifs enriched in the union of the non-differentiated chromatin accessible sites comparing ILC to IDC. **(D)** Table of the top enriched motifs in the non-differentiated chromatin accessible sites comparing the ILC and IDC chromatin accessible sites. **(E)** Table of the top enriched motifs in the chromatin accessible sites gained in IDC compared to ILC. (F) Western blot for FOXA1 of whole cell lysates of MDAMB134 cells with and without doxycycline (DOX) induction of shFOXA1.

A

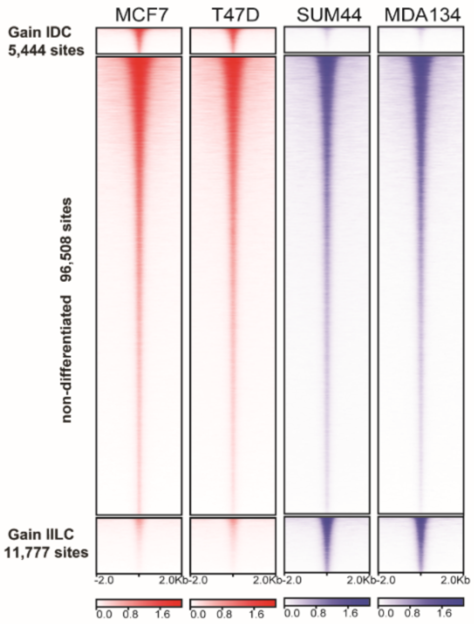


B

**Motifs Enriched in ILC gained Sites**

| MOTIF NAME | pValue | % of Target Sequences with Motif | % of Background Sequences with Motif |
|---|---|---|---|
| FOXA1(Forkhead)/ LNCAP-FOXA1-ChIP-Seq(GSE27824)/Homer | 1e-867 | 41.14% | 14.38% |
| FOXA1(Forkhead)/ MCF7-FOXA1-ChIP-Seq(GSE26831)/Homer | 1e-795 | 35.40% | 11.52% |
| FOXM1(Forkhead)/ MCF7-FOXM1-ChIP-Seq(GSE72977)/Homer | 1e-765 | 36.45% | 12.49% |
| Foxa2(Forkhead)/ Liver-Foxa2-ChIP-Seq(GSE25694)/Homer | 1e-688 | 31.96% | 10.54% |
| Fox:Ebox(Forkhead,bHLH)/ Panc1-Foxa2-ChIP-Seq(GSE47459)/Homer | 1e-520 | 38.78% | 17.42% |
| AP-2gamma(AP2)/ MCF7-TFAP2C-ChIP-Seq(GSE21234)/Homer | 1e-515 | 35.85% | 15.37% |
| Foxa3(Forkhead)/ Liver-Foxa3-ChIP-Seq(GSE77670)/Homer | 1e-515 | 44.89% | 22.22% |

C

**Motifs Enriched in non-differentiated Sites**

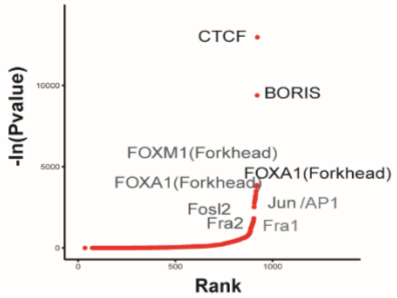


D

**Motifs Enriched in non-differentiated Sites**

| MOTIF NAME | pValue | % of Target Sequences with Motif | % of Background Sequences with Motif |
|---|---|---|---|
| CTCF(Zf)/ CD4+CTCF-ChIP-Seq(Barski_et_al.)/Homer | 1e-5630 | 14.89% | 2.20% |
| BORIS(Zf)/ K562-CTCFL-ChIP-Seq(GSE32465)/Homer | 1e-4079 | 17.70% | 4.48% |
| FOXM1(Forkhead)/ MCF7-FOXM1-ChIP-Seq(GSE72977)/Homer | 1e-1679 | 24.80% | 13.10% |
| FOXA1(Forkhead)/ MCF7-FOXA1-ChIP-Seq(GSE26831)/Homer | 1e-1667 | 24.06% | 12.57% |
| FOXA1(Forkhead)/ LNCAP-FOXA1-ChIP-Seq(GSE27824)/Homer | 1e-1641 | 27.60% | 15.38% |
| Jun-AP1(bZIP)/ K562-cJun-ChIP-Seq(GSE31477)/Homer | 1e-1524 | 8.99% | 2.80% |
| Fosl2(bZIP)/ 3T3L1-Fosl2-ChIP-Seq(GSE56872)/Homer | 1e-1517 | 11.12% | 4.02% |

E

**Motifs Enriched in IDC gained Sites**

| MOTIF NAME | pValue | % of Target Sequences with Motif | % of Background Sequences with Motif |
|---|---|---|---|
| CTCF(Zf)/ CD4+-CTCF-ChIP-Seq(Barski_et_al.)/Homer | 1e-621 | 24.50% | 2.63% |
| BORIS(Zf)/ K562-CTCFL-ChIP-Seq(GSE32465)/Homer | 1e-501 | 29.32% | 5.49% |
| GRHL2(CP2)/ HBE-GRHL2-ChIP-Seq(GSE46194)/Homer | 1.00E-43 | 9.99% | 4.67% |
| BATF(bZIP)/ Th17-BATF-ChIP-Seq(GSE39756)/Homer | 1.00E-35 | 13.69% | 7.90% |
| Atf3(bZIP)/ GBM-ATF3-ChIP-Seq(GSE33912)/Homer | 1.00E-33 | 13.82% | 8.11% |
| Fra2(bZIP)/ Striatum-Fra2-ChIP-Seq(GSE43429)/Homer | 1.00E-33 | 11.01% | 5.97% |
| JunB(bZIP)/ DendriticCells-Junb-ChIP-Seq(GSE36099)/Homer | 1.00E-33 | 12.13% | 6.83% |

F

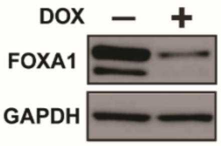


**Supplemental Figure1**

**Figure S2. FOXA1 ChIP-seq in invasive ductal cancer and invasive lobular cancer models.** **(A)** Distribution of all FOXA1 peaks. Promoters are defined as transcription factor binding sites within 1000 bp upstream of the transcription start site. Number of peaks are reported as percentages within each region. **(B)** Table of the top motifs enriched in the FOXA1 ChIP-seq sites gained in invasive lobular cancer (ILC) versus invasive ductal cancer (IDC) cells. (**C)** Table of the top motifs enriched in the union of the FOXA1 ChIP-seq sites non-differentiated between invasive lobular cancer (ILC) versus invasive ductal cancer (IDC) cell models. **(D)** Correlation of binding intensity of the sites that have gained FOXA1 binding and increased chromatin accessibility in SUM44 versus T47D cells. Spearman correlation and p-value are shown.

A

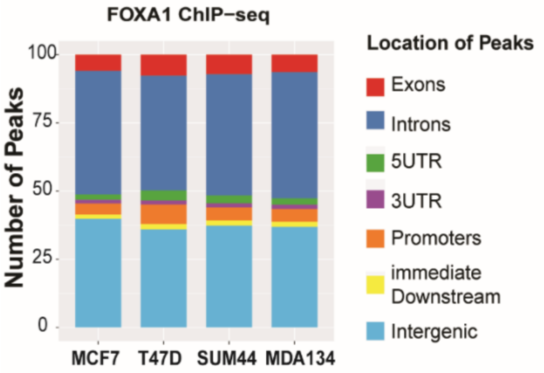


B

**Motifs Enriched in ILC gained Sites**

| MOTIF NAME | pValue | % of Target Sequences with Motif | % of Background Sequences with Motif |
|---|---|---|---|
| Foxa2(Forkhead)/ Liver-Foxa2-ChIP-Seq(GSE25694)/Homer | 1e-1340 | 57.71% | 21.72% |
| FOXA1(Forkhead)/ LNCAP-FOXA1-ChIP-Seq(GSE27824)/Homer | 1e-1297 | 70.89% | 33.32% |
| FOXA1(Forkhead)/ MCF7-FOXA1-ChIP-Seq(GSE26831)/Homer | 1e-1279 | 65.21% | 28.46% |
| FOXM1(Forkhead)/ MCF7-FOXM1-ChIP-Seq(GSE72977)/Homer | 1e-1224 | 64.63% | 28.66% |
| Foxa3(Forkhead)/ Liver-Foxa3-ChIP-Seq(GSE77670)/Homer | 1e-1055 | 34.97% | 9.56% |
| Fox:Ebox(Forkhead,bHLH)/ Panc1-Foxa2-ChIP-Seq(GSE47459)/Homer | 1e-996 | 54.81% | 23.54% |
| FoxL2(Forkhead)/ Ovary-FoxL2-ChIP-Seq(GSE60858)/Homer | 1e-766 | 52.65% | 25.07% |

C

**Motifs Enriched in non-differentiated Sites**

| MOTIF NAME | pValue | % of Target Sequences with Motif | % of Background Sequences with Motif |
|---|---|---|---|
| Foxa2(Forkhead)/ Liver-Foxa2-ChIP-Seq(GSE25694)/Homer | 1e-9587 | 54.74% | 22.75% |
| Foxa3(Forkhead)/ Liver-Foxa3-ChIP-Seq(GSE77670)/Homer | 1e-9449 | 35.70% | 10.10% |
| FOXA1(Forkhead)/ LNCAP-FOXA1-ChIP-Seq(GSE27824)/Homer | 1e-7835 | 66.93% | 36.13% |
| FOXA1(Forkhead)/ MCF7-FOXA1-ChIP-Seq(GSE26831)/Homer | 1e-7808 | 61.32% | 31.00% |
| FOXM1(Forkhead)/ MCF7-FOXM1-ChIP-Seq(GSE72977)/Homer | 1e-7352 | 60.01% | 30.65% |
| Fox:Ebox(Forkhead,bHLH)/ Panc1-Foxa2-ChIP-Seq(GSE47459)/Homer | 1e-6373 | 48.97% | 23.10% |
| FOXK2(Forkhead)/ U2OS-FOXK2-ChIP-Seq(E-MTAB-2204)/Homer | 1e-5498 | 39.83% | 17.55% |

D

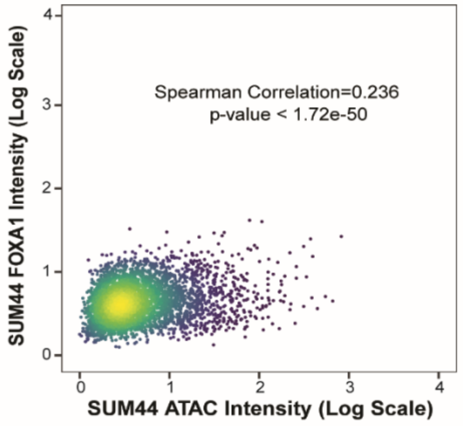


**Supplemental Figure 2**

**Figure S3. ER binding and H3K27ac ChIP-seq profiles of IDC and ILC models**

**(A)** Distribution of ER peaks. Promoters are defined as TF binding sites within 1000 bp upstream of the TSS. Number of peaks are reported as percentages in each region. **(B)** Table of top motifs enriched in ER binding sites gained in invasive lobular cancer (ILC) compared to invasive ductal cancer (IDC) cells. **(C)** Table of top motifs enriched in the union of the ER binding sites non-differentiated between ILC IDC cells. **(D)** Sample to sample correlation plot of H3K27ac sites in ILC and IDC cells. **(E)** Tornado plot of all H3K27ac binding events in the four cell line models and primary cells from an ILC estrogen receptor positive metastatic peritoneal effusion.

A

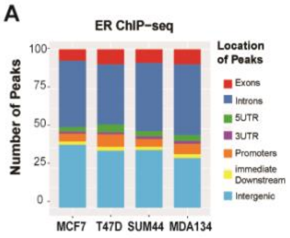


B

**Motifs Enriched in ILC gained Sites**

| MOTIF NAME | pValue | % of Target Sequences with Motif | % of Background Sequences with Motif |
|---|---|---|---|
| ERE(NR),IR3/ MCF7-ERa-ChIP-Seq(Unpublished)/Homer | 1e-938 | 34.65% | 6.34% |
| FOXA1(Forkhead)/ LNCAP-FOXA1-ChIP-Seq(GSE27824)/Homer | 1e-271 | 40.17% | 20.38% |
| FOXA1(Forkhead)/ MCF7-FOXA1-ChIP-Seq(GSE26831)/Homer | 1e-269 | 35.14% | 16.56% |
| COUP-TFII(NR)/ Artia-Nr2f2-ChIP-Seq(GSE46497)/Homer | 1e-262 | 59.88% | 37.91% |
| FOXM1(Forkhead)/ MCF7-FOXM1-ChIP-Seq(GSE72977)/Homer | 1e-261 | 36.16% | 17.57% |
| Foxa2(Forkhead)/ Liver-Foxa2-ChIP-Seq(GSE25694)/Homer | 1e-229 | 30.78% | 14.47% |
| Fox:Ebox(Forkhead,bHLH)/ Panc1-Foxa2-ChIP-Seq(GSE47459)/Homer | 1e-199 | 36.29% | 19.66% |

C

**Motifs in non-differentiated Sites**

| MOTIF NAME | pValue | % of Target Sequences with Motif | % of Background Sequences with Motif |
|---|---|---|---|
| ERE(NR),IR3/ MCF7-ERa-ChIP-Seq(Unpublished)/Homer | 1e-2940 | 20.09% | 4.96% |
| COUP-TFII(NR)/ Artia-Nr2f2-ChIP-Seq(GSE46497)/Homer | 1e-1418 | 49.76% | 31.87% |
| FOXA1(Forkhead)/ LNCAP-FOXA1-ChIP-Seq(GSE27824)/Homer | 1e-1197 | 44.49% | 28.47% |
| FOXM1(Forkhead)/ MCF7-FOXM1-ChIP-Seq(GSE72977)/Homer | 1e-1179 | 39.76% | 24.43% |
| FOXA1(Forkhead)/ MCF7-FOXA1-ChIP-Seq(GSE26831)/Homer | 1e-1167 | 39.33% | 24.13% |
| Foxa2(Forkhead)/ Liver-Foxa2-ChIP-Seq(GSE25694)/Homer | 1e-1062 | 32.20% | 18.76% |
| Fox:Ebox(Forkhead,bHLH)/ Panc1-Foxa2-ChIP-Seq(GSE47459)/Homer | 1e-943 | 34.27% | 21.17% |

D

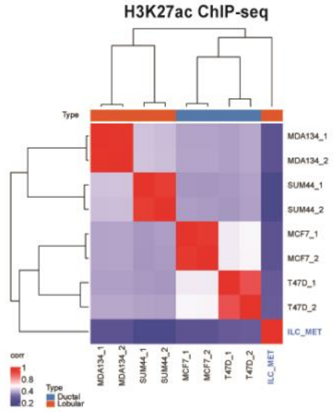


E

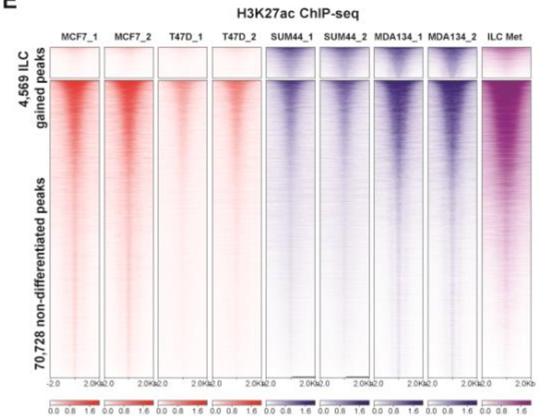


F

**Motifs Enriched in ILC gained Sites**

| MOTIF NAME | pValue | % of Target Sequences with Motif | % of Background Sequences with Motif |
|---|---|---|---|
| FOXA1(Forkhead)/ LNCAP-FOXA1-ChIP-Seq(GSE27824)/Homer | 1.e-115 | 36.89% | 21.04% |
| FOXM1(Forkhead)/ MCF7-FOXM1-ChIP-Seq(GSE72977)/Homer | 1e-105 | 32.67% | 18.19% |
| FOXA1(Forkhead)/ MCF7-FOXA1-ChIP-Seq(GSE26831)/Homer | 1e-104 | 31.37% | 17.22% |
| Foxa2(Forkhead)/ Liver-Foxa2-ChIP-Seq(GSE25694)/Homer | 1e-81 | 26.40% | 14.70% |
| Fox:Ebox(Forkhead,bHLH)/ Panc1-Foxa2-ChIP-Seq(GSE47459)/Homer | 1e-74 | 31.92% | 19.61% |
| Foxa3(Forkhead)/ Liver-Foxa3-ChIP-Seq(GSE77670)/Homer | 1e-67 | 13.09% | 5.68% |
| PHA-4(Forkhead)/cElegans-Embryos -PHA4-ChIP-Seq(modEncode)/Homer | 1e-62 | 64.97% | 51.83% |

G

**Motifs in non-differentiated Sites**

| MOTIF NAME | pValue | % of Target Sequences with Motif | % of Background Sequences with Motif |
|---|---|---|---|
| FOXM1(Forkhead)/ MCF7-FOXM1-ChIP-Seq(GSE72977)/Homer | 1e-440 | 28.23% | 20.44% |
| FOXA1(Forkhead)/ LNCAP-FOXA1-ChIP-Seq(GSE27824)/Homer | 1e-438 | 31.91% | 23.76% |
| FOXA1(Forkhead)/ MCF7-FOXA1-ChIP-Seq(GSE26831)/Homer | 1e-431 | 27.70% | 20.03% |
| Foxa2(Forkhead)/ Liver-Foxa2-ChIP-Seq(GSE25694)/Homer | 1e-383 | 22.39% | 15.76% |
| Foxa3(Forkhead)/ Liver-Foxa3-ChIP-Seq(GSE77670)/Homer | 1e-317 | 10.80% | 6.56% |
| Fox:Ebox(Forkhead,bHLH)/ Panc1-Foxa2-ChIP-Seq(GSE47459)/Homer | 1e-317 | 24.89% | 18.60% |
| PHA-4(Forkhead)/cElegans-Embryos -PHA4-ChIP-Seq(modEncode)/Homer | 1e-233 | 58.97% | 52.24% |

**Supplemental Figure 3**

**Figure S4. FOXA1 transcriptional target genes upregulated in invasive lobular cancer**
**(A)** Binding and Expression Target Analysis **(**BETA) basic plot of the activating and repressive function of the FOXA1 binding sites gained in invasive lobular cancer (ILC) cells compared to invasive ductal (IDC) cells. The red line represents the genes upregulated and the purple line the genes downregulated comparing ILC versus IDC RNASeq. The black dashed line indicates the non-differentially expressed genes as background. The p-value is based on the Kolmogorov-Smirnov test. **(B)** Volcano plot depicting differential (DEseq2) expression comparing RNA-seq of MDAMB134 versus MCF7 cells after β-estradiol stimulation (E2). Shown in yellow are the genes with significant differential expression ([log2FC] >1, *Q*<0.01. DESeq2). Red dots represent the genes that are significantly differentially expressed ([log2FC] >1, *Q*<0.01. DESeq2) and are regulated by FOXA1 ILC gained binding sites based on Binding and Expression Target (BETA) minus analysis. (**C)** Volcano plot showing the results of differential gene expression of MDAMB134 cells with and without silencing of FOXA1. **(D)** Binding and Expression Target Analysis **(**BETA) basic plot of the activating and repressive function of the FOXA1 binding sites gained in invasive lobular cancer (ILC) cells compared to invasive ductal (IDC) cells and the genes regulated by FOXA1. The red line represents the genes upregulated and the purple line the genes downregulated comparing RNA-seq of MDAMB134 cells with and without FOXA1 silencing. The black dashed line indicates the non-differentially expressed genes as background. The p-value is based on the Kolmogorov-Smirnov test. **(E)** ChIP-seq tracks showing ER, FOXA1, and H3K27acetylation in the genomic regions of *RET, SNAI1*, and *CASZ1* in ILC (MDA134 (MDAMB134) and SUM44, in purple and blue respectively) and IDC (MCF7 and T47D in red and orange respectively) cells. (**F**) Western blot of whole cell lysates of MCF7 and MDA134 cells transfected with siControl (CTR), siER or siFOXA1. Numbers represent protein quantification. **(G)** CASZ1 mRNA expression in the METABRIC cohort comparing invasive ductal and lobular

breast cancers. **(H)** Cell proliferation studies of MDA134 cells in control cells (siCTR) and cells with silencing of CASZ1 (siCASZ1_1 and siCASZ1_2). Error bars represent ±SEM, n=3. **(I)** Genes differentially expressed after siCASZ1 (FDR<0.05) in MDA134 cells compared to siCTR. *=p<0.05; **= p<0.01, Student's T-test. **(J)** Immunoblotting of whole cell lysates for CASZ1, CDK2, FOXA1, ER and GAPDH in siCTR and siCASZ1 in MDA134 cells.

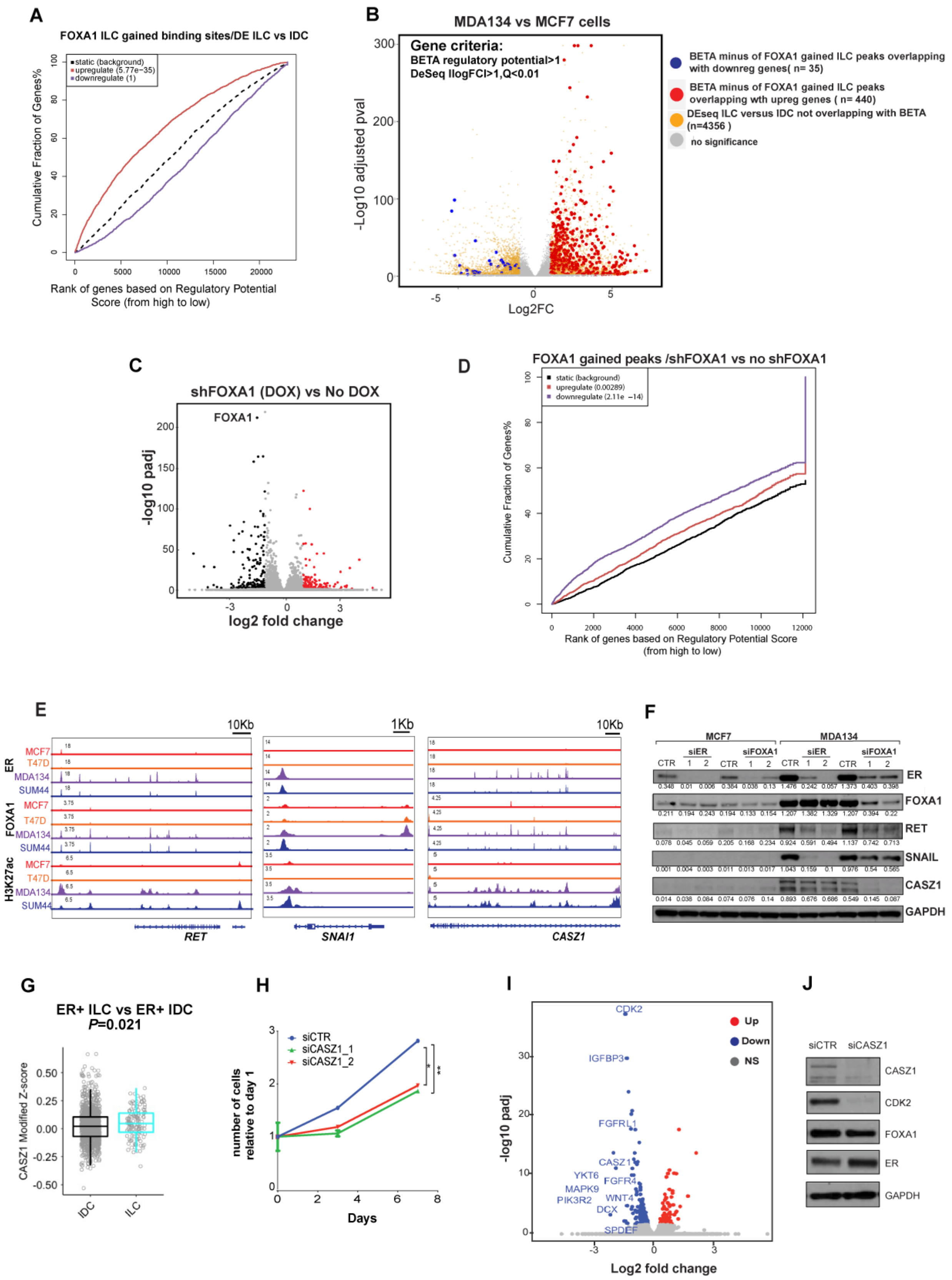


**Supplemental Figure 4**

**Figure S5. Transcriptional effects of E-cadherin silencing and overexpression in estrogen receptor positive breast cancer cells.**

**(A)** Western blotting for E-cadherin of whole cell lysates of MCF7, T47D, MDAMB134 (MDA134) and SUM44 cells, and MCF7 and T47D induced with a control gRNA (gAAVS1) or a gRNA targeting *CDH1* (gCDH1) for down regulation of E-cadherin expression. **(B)** Immunofluorescence staining for E-cadherin (green) and nuclear staining with DAPI (blue). **(C)** MCF7 and T47D were subjected to trans-well migration assays for 48hrs. Bar graph shows the number of migrated cells in control (gAAVS1) and cells with E-cadherin down regulation (gCDH1). Error bars represent mean ± SEM. *=p-value < 0.05, *** =p-value < 0.00, Student's T-test. **(D)** Cell proliferation studies in MCF7 and T47D in control cells (gAAVS1) and cells with E-cadherin down regulation (gCDH1). Error bars represent ±SEM, n = 3. **(E)** Volcano plot of genes differentially expressed in MCF7 cells with and without E-cadherin down regulation (FDR<0.05). **(F)** Pathways enriched in the genes differentially expressed with downregulation of E-cadherin in MCF7 cells based on GSEA (q-value <0.25)**.** **(G)** Volcano plot of genes differentially expressed in T47D cells with and without E-cadherin down regulation (FDR<0.05). **(H)** Pathways enriched in the genes differentially expressed with downregulation of E-cadherin in T47D cells based on GSEA (q-value <0.25)**. (I)** Tornado plots of the ER binding events gained in lobular cells compared with ductal cells. ER binding sites are shown in a horizontal window of ±2 kb from the peak center. **(J)** Tornado plots of the H3K27 acetylation gained in lobular cells compared with ductal cells. H3K27acetylation is shown in a horizontal window of ±2 kb from the peak center. **(K-L)** Volcano plots of differential genes in MCF7 (K) and T47D (L) cells with and without down regulation of E-cadherin in estrogen (E2) stimulated conditions. (M) Western blot of whole cell lysates from MDAMB134 cells stably expressing doxycycline (DOX) induced E-cadherin. E-cadherin expression is shown in cells not treated with DOX and cells treated with

DOX for 2 days and 3 weeks. **(N-O)** Volcano plots of differential genes in MDAMB134 cells with and without DOX induction of E-cadherin expression at 2 days **(N)** and 3 weeks **(O)** of E-cadherin expression.

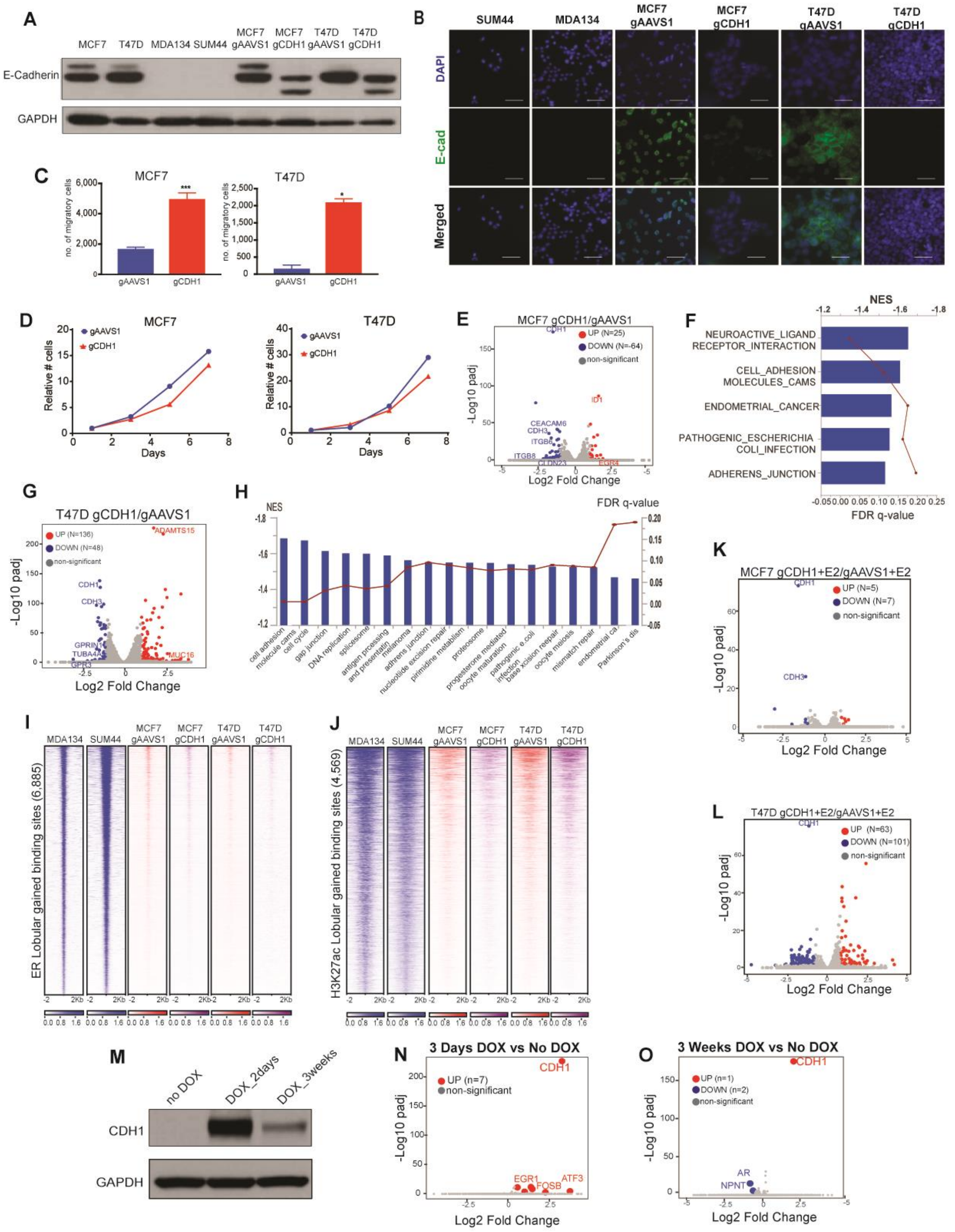


**Supplemental Figure 5**

**Figure S6. Genomic Landscape and H3K27ac Hi-ChIP looping**

**(A)** Oncoprint of single nucleotide variants of 72 breast cancer related genes identified using the GATK for SNP and INDEL. The histogram on the right shows the frequency of each alteration in each cell line. Lower panel: Copy number variations (CNVs) of breast cancer related genes detected in the same gene panel using whole genome sequencing. Red indicates copy number gain and blue copy number loss. **(B)** H3K27ac-HiChIP-seq tracks showing looping at the FOXA1 promoter regions in the invasive ductal cancer cell lines, MCF7 and T47D in red, and the invasive lobular cancer cells, MDAMB134 (MDA134) and SUM44 in blue. **(C)** Cell proliferation study in MCF7 cells transfected with siControl (siCTR) and siFOXA1. Error bars represent ±SEM, n = 3. ****, p-value <0.001(Student's t-test) **(D)** Flow cytometry analysis for GFP (left) and BFP (right) in control MCF7 cells, MCF7 cells expressing the GFP/BFP reporter plasmid with a gRNA targeting GFP without dCAS9-KRAB or CAS9 and MCF7 cells expressing dCAS9-KRAB and the GFP/BFP reporter plasmid with a gRNA targeting GFP.

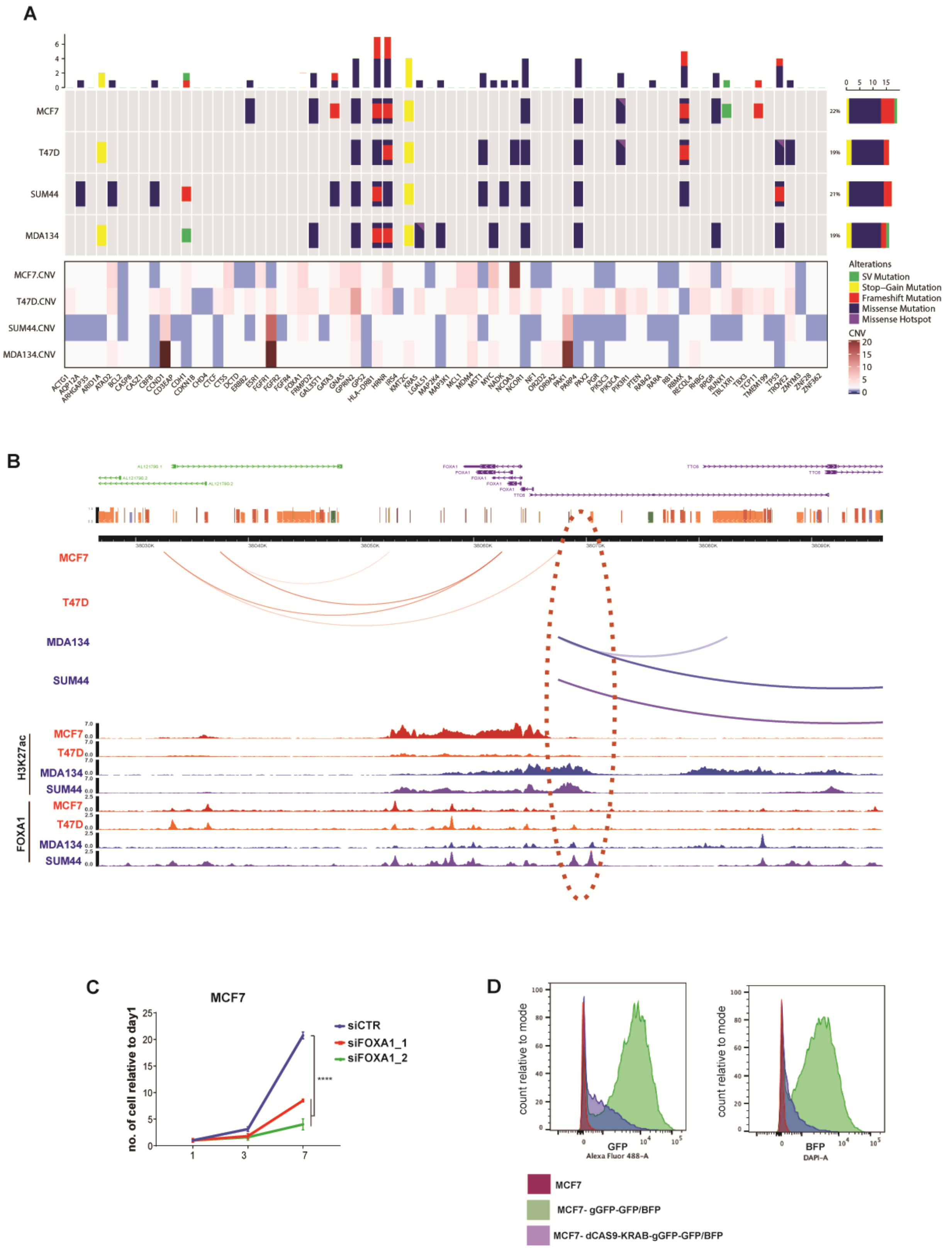


**Supplemental Figure 6**

**Figure S7. ILC cells are resistant to tamoxifen but responsive to ED, siER and siFOXA1**

**(A)** Cell proliferation studies of MCF7 cells followed for 7 days in hormone deprived conditions (HD) without estrogen or with estrogen (estradiol (E2) 10nM) for 7 days. Error bars represent ±SEM, n = 3. ****= p value <0.0001, Student's T-test. **(B)** Cell proliferation studies of MCF7 cells in full medium conditions including cells transfected with an siControl (siCTR) and cells with silencing of ER (siER_1 and siER_2). Error bars represent ±SEM, n = 3. ****= p value <0.0001, Student's T-test**.** **(C)** Number of ER binding that are either gained or lost with 4-hydroxytamoxifen treatment (TAM) compared to estrogen (E2,10nM) treatment in MCF7 cells and MDAMB134 (MDA134) cells. D. Normalized signal enrichment of chromatin accessible sites at the ER binding sites that were lost in MCF 7 cells but unchanged in MDAMB134 cells (MDA134). E. Cell proliferation studies of MDAMB134 cells with doxycycline (DOX) or without doxycycline (ND) induction of shFOXA1. Error bars represent ±SEM, n = 3. **= p value <0.001, Student's T-test**.**

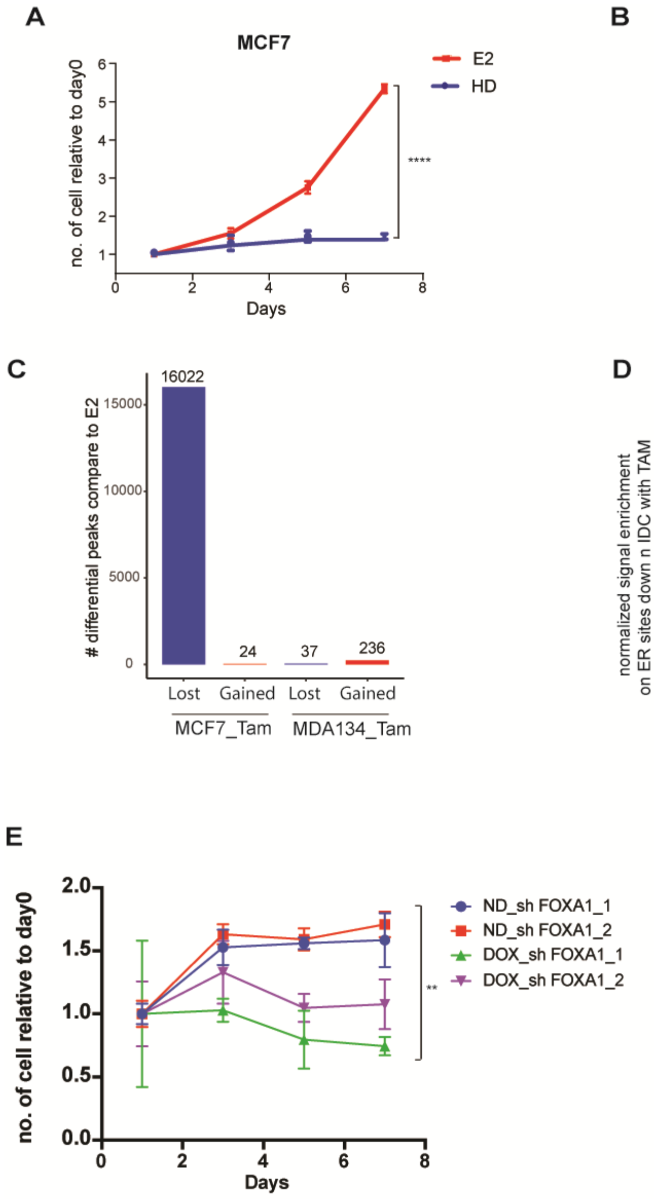


**Supplemental Figure 7**
